# Nuclear autophagy interactome unveils WSTF as a constitutive nuclear inhibitor of inflammation

**DOI:** 10.1101/2022.10.04.510822

**Authors:** Yu Wang, Vinay V. Eapen, Athanasios Kournoutis, Angelique Onorati, Xianting Li, Xiaoting Zhou, Murat Cetinbas, Lu Wang, Jihe Liu, Corey Bretz, Zhuo Zhou, Shannan J. Ho Sui, Srinivas Vinod Saladi, Ruslan I. Sadreyev, Peter D. Adams, Robert E. Kingston, Zhenyu Yue, Terje Johansen, Zhixun Dou

## Abstract

Macroautophagy (hereafter referred to as autophagy) degrades a variety of cellular components. A poorly understood area is autophagic degradation of nuclear substrates, or “nuclear autophagy”. It remains unclear what can be degraded by autophagy from the mammalian nuclei. We began our study by investigating the nuclear binding partners of ATG8 family proteins that play important roles in recognizing autophagy substrates. We systematically evaluated the ATG8 nuclear interactome in primary human cells and in mouse brain, identifying hundreds of novel interactions. We continued our study by evaluating the nuclear proteomes of cellular senescence, a stable form of cell cycle arrest program associated with inflammation, in which nuclear autophagy is involved. Combined with the ATG8 nuclear interactome data, we identified WSTF, a component of the ISWI chromatin remodeling complex, as a novel substrate of nuclear autophagy. The degradation of WSTF, mediated by a direct interaction with the GABARAP isoform of ATG8, promotes chromatin accessibility of inflammatory genes and induces senescence-associated inflammation. Furthermore, WSTF directly binds the p65 subunit of NF-κB and inhibits its acetylation, thus blocking inflammatory gene expression in the setting of senescence, cancer, and pathogen infection. In addition, we show that loss of WSTF is required for the immuno-surveillance of oncogenic Ras in mouse liver; forced expression of WSTF inhibited tumor-suppressive inflammation and led to the development of liver tumors. Taken together, our study provides a global view of mammalian nuclear autophagy and reveals a novel nuclear inhibitor of inflammation implicated in diverse pathological contexts. Targeting WSTF may be broadly valuable as therapeutic intervention of inflammatory diseases.

## Introduction

Mammalian ATG8 family proteins (including LC3A, LC3B, LC3C, GABARAP, GABARAPL1, and GABARAPL2) bind to selective autophagy receptors or directly to autophagy substrates and help mediate their delivery to autophagosomes for degradation^1^. While yeasts have one single ATG8 protein, the six isoforms of mammalian ATG8 family proteins have both distinct and redundant roles in autophagosome formation, expansion, maturation, vesicle transport, docking of autophagy receptors, and in direct recognition of autophagy substrates^2, 3^. ATG8 proteins are present in the nucleus with low mobilities^4, 5^, suggesting that they interact with large molecular weight complexes. We recently reported that nuclear ATG8s directly bind to nuclear Lamin B1 and SIRT1, facilitating their shuttling to the cytoplasm for autophagic degradation upon cellular senescence^6, 7^, suggesting that autophagy has a previously unappreciated role in degrading nuclear constituents, termed as “nuclear autophagy”. Beyond Lamin B1 and SIRT1, whether nuclear autophagy can degrade other substrates is unclear, which represents a major open area for investigation.

Cellular senescence is a pro-inflammatory condition in which nuclear autophagy is induced^6, 7^. Senescent cells accumulate in aged and diseased tissues, where they impair tissue renewal and promote inflammation^8, 9^. While senescence is a potent tumor-suppressive mechanism, it paradoxically contributes to aging and the progression of many diseases^10–12^. Clearing senescent cells using pharmacological or genetic approaches ameliorates several diseases in mice and humans^13, 14^. A key feature of senescence is the secretion of a large array of pro-inflammatory cytokines, chemokines, growth factors, and proteases, collectively referred to as senescence-associated secretory phenotype (SASP)^8, 9, 11, 12^. The SASP program recruits immune cells and alters tissue microenvironment, leading to inflammation^8, 9, 11, 12^. Targeting the SASP program is an important biomedical objective to intervene in diseases. The roles of nuclear autophagy in senescence are poorly understood, largely due to our limited knowledge of the substrates of nuclear autophagy.

To fill the major gaps in our understanding of nuclear autophagy, we examined the nuclear binding partners of ATG8s as well as the nuclear proteome of senescent cells. These proteomic studies offer a comprehensive resource to study nuclear autophagy. We further focused on WSTF, which we identified as a new substrate of nuclear autophagy, and discovered an unexpected role of WSTF in regulating inflammation.

## Results

### The nuclear ATG8 interactome

While ATG8 interactomes using whole cell extracts have been reported by several groups^15–17^, very few nuclear interactions have been identified. We reason that this is likely due to the poor lysis efficiency of the nuclear and chromatin fraction. Most interactomics studies used NP40 or Triton X-100 to lyse cells; after a centrifugation step, the supernatants were collected for subsequent co-immunoprecipitation (co-IP). These standard lysis conditions led the chromatin fraction to become pelleted and discarded for the subsequent IP steps. Using histone H3 as a marker for the chromatin fraction, we found that 1% NP40 lysis showed a minimal signal of H3 in the supernatant (Fig. 1a). To resolve the poor solubility issue of the chromatin fraction, we used benzonase, an endonuclease that digests chromatin DNA, to facilitate the release of chromatin proteins to the supernatant. The efficiency is similar to that of adding 1% SDS followed by sonication (Fig. 1a). Thus, we generated a new procedure suitable for nuclear interactome studies.

**Figure 1.**
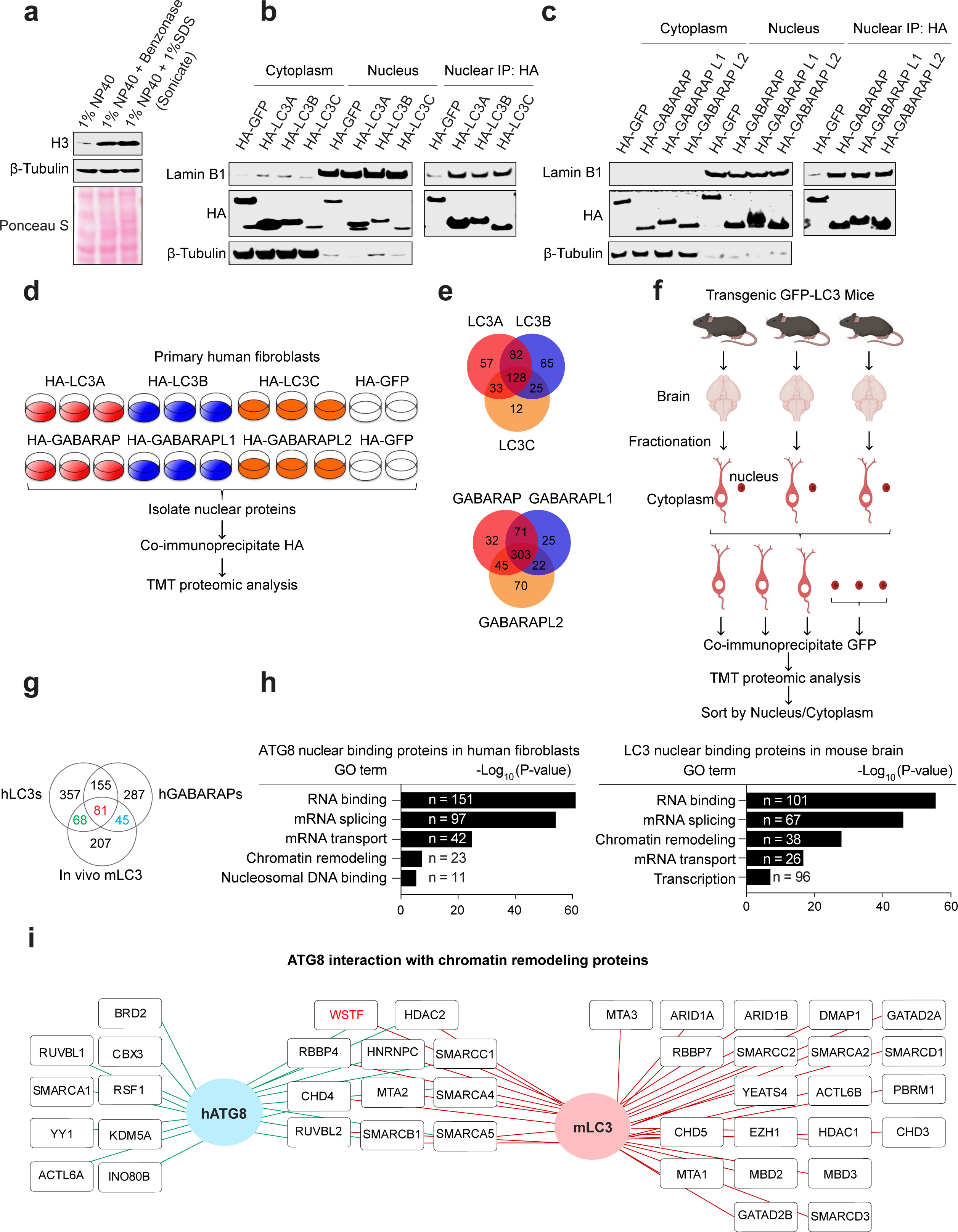
Generation of nuclear ATG8 interaction network. **a**, Primary IMR90 cells were subjected to protein extraction using different approaches. Supernatants were blotted with indicated antibodies. **b, c,** Primary IMR90 cells cultured in 3% oxygen were stably infected by lentiviral constructs encoding HA-tagged GFP or ATG8 genes. The cells were subjected to subcellular fractionation, and the nuclear and chromatin fractions (collectively referred to as “nucleus”) were subjected to co-immunoprecipitation using anti-HA antibody. The input and IP products were analyzed by immunoblotting. **d,** Scheme depicting the workflow of TMT proteomic analysis, using cells as described in **b** and **c**. **e,** ATG8 nuclear interactomes were analyzed and targets with P values less than 0.05 (compared to GFP negative control) and log2 ATG8/GFP > 0 were included in the Venn diagrams. The numbers of nuclear binding partners of each ATG8 member, and the overlap of LC3 or GABARAP subgroups are shown. **f,** Scheme depicting the workflow of TMT proteomic studies using the brain of GFP-LC3B transgenic mice. WT (non-transgenic) mice were also included in these experiments (not shown in the scheme). **g,** Venn diagram showing the numbers of nuclear binding partners of human LC3A, LC3B and LC3C (hLC3s), and human GABARAP, GABARAPL1 and GABARAPL2 (hGABARAPs), and LC3B from mouse brain. **h,** Gene Ontology (GO) analyses of ATG8 nuclear binding proteins in human fibroblasts and LC3B nuclear binding proteins in mouse brain. Top biological processes, together with protein numbers and P values, are shown. **I,** Protein-protein interaction map presentation of ATG8 nuclear binding partners in human fibroblasts and mouse brain, focusing on proteins involved in chromatin remodeling.

We then stably expressed the six ATG8 isoforms in IMR90 primary human fibroblasts, cultured in physiological 3% oxygen, and performed nuclear fractionation and co-IPs (Fig. 1b and 1c). As a positive control, an interaction with Lamin B1 was detected (Fig. 1b and 1c). We subsequently performed tandem mass tag (TMT)-based mass-spectrometry analyses, with three biological replicates of each ATG8 isoform and two biological replicates of GFP as negative controls (scheme shown in Fig. 1d). A total of 422 statistically significant nuclear interactions were identified for LC3A, LC3B, LC3C, and 568 for GABARAP, GABARAPL1, GABARAPL2 (Fig. 1e). As expected, these isoforms showed overlapping yet distinct binding partners (Fig. 1e). Top interaction partners are presented in interaction maps (Extended data Fig. 1a and 1b). When comparing with a previous autophagy interaction network (AIN) focusing on cytoplasmic interactions^15^, we found that the majority of the binding partners of ATG8 from our nuclear AIN are unique and previously unknown (Extended data Fig. 1c). Hence, we generated a new database of nuclear ATG8 interactome in primary human cells.

To further explore ATG8 binding partners *in vivo*, we used a GFP-LC3B transgenic mouse strain^18^ and performed GFP co-IPs in mouse brain, in both cytoplasmic and nuclear fractions (scheme shown in Fig. 1f). Wild-type (not expressing GFP-LC3B) mice were used as negative controls. Similar to the nuclear fractionation of primary human cells, a benzonase treatment step was included, leading to successful solubilization of histone H3 (Extended data Fig. 1d). Compared with the negative control using wild-type mice, GFP co-IPs in GFP-LC3B mice precipitated specific proteins (Extended data Fig. 1e). Three biological replicates for cytoplasmic GFP co-IPs were included, and due to the low protein concentrations of the nuclear fractions, the three nuclear fractions were combined for TMT-based mass-spectrometry analyses (Fig. 1f). Although LC3B was present much less in the nucleus than in the cytoplasm, specific binding with nuclear proteins was detected from nuclear GFP co-IPs.

We went on further to analyze the nuclear binding partners of ATG8s in primary human cells and mouse brain, and discovered the overlapping and unique interactions in the two biological models (Fig. 1g and specific proteins listed in Extended data Fig. 1g). Gene ontology (GO) analyses revealed similar profiles of the nuclear binding partners of ATG8s in the two models, with substantial proteins within the categories of RNA binding, mRNA splicing and transport (Fig. 1h). The binding between ATG8s and RNA-related proteins is supported by recent studies showing LC3 mediates RNA secretion^19^ and degradation^20^. In addition, proteins involved in chromatin remodeling, nucleosomal binding, and transcription were identified (Fig. 1h). The overlapping nuclear proteins from the two unrelated biological models suggest conserved roles for nuclear ATG8s across tissues in mammals in regulating these biological processes. Of our particular interest is the chromatin remodeling complexes that bind to both human and mouse ATG8s (Fig. 1i), as these proteins regulate nucleosome positioning and chromatin structures, thereby affecting gene expression and silencing.

Taken together, we have generated a systematic resource of nuclear ATG8 interactomes in primary human cells and mouse brain, and identified chromatin remodeling complexes as putative ATG8 binding partners in mammalian nuclei.

### Nuclear proteomes of cellular senescence

We previously reported that nuclear Lamin B1 and SIRT1 autophagic degradation occurs in cellular senescence, in which cells respond to cellular insults such as oncogene activation and telomere attrition and upregulate autophagy activities^6, 7^. To explore novel substrates of nuclear autophagy under senescence, we induced senescence of primary IMR90 cells using two different means, oncogene activation and sub-lethal dose of DNA damage, and searched for overlapping hits with ATG8 nuclear interactomes.

We began by investigating the whole cell proteome of HRasV12-induced senescence (Fig. 2a), using TMT-based mass-spectrometry, done with three biological replicates (Fig. 2b and Extended data Fig. 2a). Consistent with our existing knowledge of senescence, the proteome of senescent cells harbors significantly upregulated proteins, most of which are involved in the SASP program and the DNA damage responses, as well as significantly downregulated proteins involved in cell cycle and DNA replication (Fig. 2c and Extended data Fig. 2b). Among the downregulated proteins, 718 are nuclear proteins (Fig. 2d), and 89 of them (including Lamin B1) overlap with ATG8 nuclear interactome of IMR90 cells (Fig. 2e and listed in Extended data Fig. 2c).

**Figure 2.**
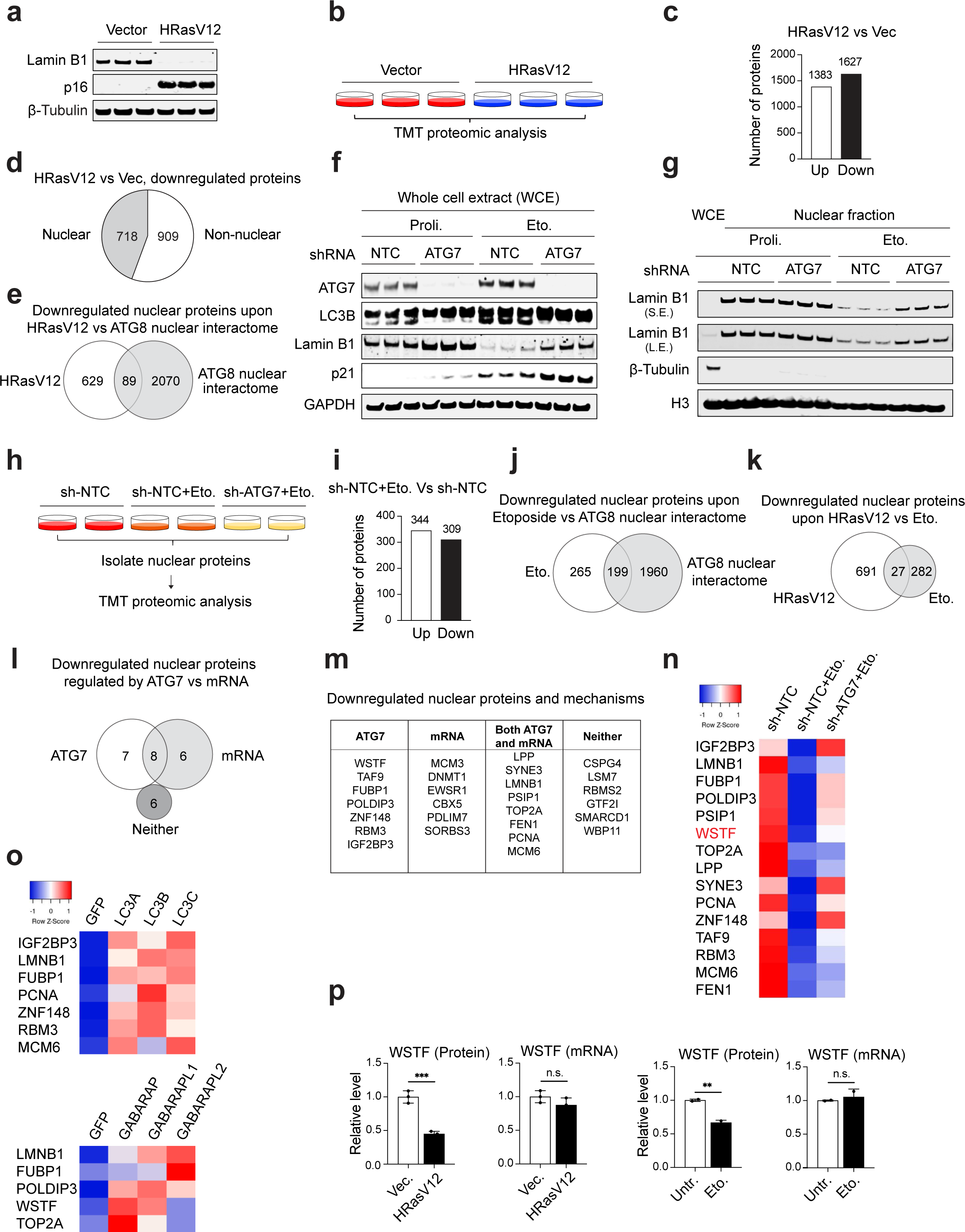
Nuclear proteome analyses of senescent cells. **a**, Primary low passage IMR90 cells cultured in 3% oxygen were infected with retrovirus encoding HRasV12 or vector control. 7 days post the infection, cell lysates were analyzed by western blotting with indicated antibodies. Three independent replicates were used for each condition. **b**, Scheme depicting the workflow for TMT proteomic analysis, using cells as described in **a**. **c**, Bar graph showing the numbers of upregulated and downregulated proteins in HRasV12 cells compared to vector control cells. For upregulated, proteins that meet log2 HRasV12/Vector > 0.5, p < 0.05 were included; for downregulated, proteins that meet log2 HRasV12/Vector < -0.5, p < 0.05 were included. **d**, Pie chart showing the numbers of nuclear and non-nuclear proteins within the downregulated proteins in HRasV12 cells from **c**, as defined by GO term cellular component. **e,** Venn diagram showing the numbers of downregulated nuclear proteins upon HRasV12 overlapping with ATG8 nuclear interactome from human fibroblasts. **f,** Primary low passage IMR90 cells were infected with lentivirus expressing non-targeting control hairpin (sh-NTC) or sh-Atg7 hairpin. The cells were treated with 50 µM etoposide for 2 days and then cultured without etoposide for 7 days. Whole cell extract (WCE) was analyzed by western blotting with antibodies shown. **g**, IMR90 cells as described in **f** were subjected to subcellular fractionation. Whole cell extract (WCE) and nuclear fraction were analyzed by western blotting with antibodies indicated. **h**, Scheme depicting the workflow of the TMT proteomic analyses, using nuclear fractions as described in **g**. **i**, Bar graph showing the numbers of upregulated and downregulated nuclear proteins in etoposide-treated cells compared to control cells. For upregulated, proteins meet log2 Eto/CT > 0.5, p < 0.05 were included; for downregulated, proteins meet log2 Eto/CT < -0.5, p < 0.05 were included. **j**, Venn diagram showing the numbers of downregulated nuclear proteins upon etoposide overlapping with ATG8 nuclear interactome from primary human fibroblasts. **k,** Venn diagram showing the numbers of downregulated nuclear proteins in HRasV12-induced senescence overlapping with etoposide-induced senescence. **l**, Venn diagram showing the numbers of decreased nuclear proteins regulated by ATG7 and/or mRNA. **m**, Related to **l**, table showing the names of decreased nuclear proteins regulated by ATG7 and/or mRNA. **n**, Heatmap presentation of relative protein levels from indicated groups. Note that sh-ATG7 rescued the protein downregulation in sh-NTC etoposide group. **o**, Proteins as in **n** were overlapped with ATG8 nuclear interactomes from human fibroblasts. The overlapped proteins were analyzed for their binding affinities with ATG8 and were presented as heatmaps. **p,** Bar graphs showing the protein and mRNA levels of WSTF upon HRasV12 and etoposide-induced senescence.

We further examined the proteome of etoposide-induced senescence, in both autophagy-competent and autophagy-deficient cells, using shRNA-mediated silencing of ATG7, focusing on nuclear fractions (Fig. 2f-2h and Extended data Fig. 2d). As expected, inhibition of ATG7 suppressed LC3 lipidation and impaired Lamin B1 downregulation in senescence (Fig. 2f and 2g). The nuclear fractions of these cells were subjected to TMT-based mass-spectrometry analyses (Fig. 2h and Extended data Fig. 2e). 309 proteins were downregulated in the autophagy competent cells and 199 of them (including Lamin B1) overlap with ATG8 nuclear interactome of IMR90 cells (listed in Extended data Fig. 2f). We did not detect SIRT1 in these proteomic studies, likely due to the higher sensitivity of SIRT1 antibodies than that of mass-spectrometry.

While HRasV12 and etoposide-induced senescence both revealed putative downregulated nuclear proteins that bind to ATG8, we reason that the 27 overlapping targets in the two senescence models (Fig. 2k) are of greater generalizability in senescence. We thus investigated these targets in greater detail, and further asked whether their downregulation is linked to ATG7 and/or their mRNA levels using our previously established RNA-seq datasets^7, 21^. We grouped these targets in four categories: (1) rescued by ATG7 deficiency while their mRNA levels are unaltered, (2) not rescued by ATG7 deficiency while their mRNA levels are reduced; (3) rescued by ATG7 deficiency while their mRNA levels are also reduced in senescence, and (4) not linked to ATG7 or mRNA downregulation (Fig. 2l and 2m). As a control, Lamin B1 is known to be downregulated at both mRNA and protein levels^6^, and is listed in group (3). The proteins that are lost in senescence and are rescued by ATG7 deficiency are further presented in a heatmap (Fig. 2n) and overlapped with ATG8 nuclear interactome (Fig. 2o).

Because our ATG8 nuclear interactome studies identified chromatin remodeling proteins as putative binding partners of ATG8 (Fig. 1h and 1i), we examined their protein levels in senescence (Extended data Fig. 2g). Our results showed that WSTF is a consistent target in all our proteomic datasets: WSTF binds to ATG8 in both human fibroblasts and mouse brain (Fig. 1i and Extended data Fig. 1b and 1g) and is downregulated in senescent cells, which can be rescued by ATG7 deficiency (Fig. 2m-2o and Extended data 2g). Further, while WSTF protein is reduced in senescence, its mRNA levels remain unchanged (Fig. 2m and 2p), making it a strong candidate as a substrate of nuclear autophagy. WSTF (also known as BAZ1B, hereafter referred to as WSTF for both gene and protein), is a subunit of imitation switch (ISWI) chromatin remodeling complex^22^. The ISWI chromatin remodeling complex regulates nucleosome spacing, changing the chromatin from an “open” state to a “closed” state, by forming ordered nucleosome arrays on chromatin, thereby repressing gene expression^23–25^. WSTF has not been investigated in the context of autophagy or senescence. In summary, we generated nuclear proteomic datasets in two forms of senescence, and revealed new potential substrates of nuclear autophagy. We will focus on WSTF for the rest of this study, investigating its autophagic degradation and its biological functions.

### WSTF is degraded by autophagy in senescence

We induced senescence by multiple means, in multiple cell systems, to investigate the behavior of WSTF. WSTF protein levels are dramatically reduced in senescent IMR90 cells induced by etoposide (Fig. 3a), HRasV12 (Fig. 3b), or replication exhaustion (Fig. 3c). The loss of WSTF protein is also observed in senescent primary BJ fibroblasts, induced by etoposide, ionizing irradiation (IR), or HRasV12, as well as in senescent primary MEFs induced by etoposide or IR (Extended data Fig. 3a). In addition, therapy-induced senescence of A549 cancer cells exhibited loss of WSTF protein (Extended data Fig. 3b). In contrast to the loss of WSTF in senescence, WSTF protein levels remain unchanged upon quiescence or starvation (Extended data Fig. 3c-3e). We further examined other proteins of the ISWI complex. SNF2H (also known as SMARCA5), the ATPase of the ISWI complex that binds to WSTF, is largely unaltered in senescence (Fig. 3a). Another subunit of the ISWI complex, ACF1 (also known as BAZ1A) that also binds to SNF2H, is reduced in senescence (Fig. 3a). However, ACF1 is also reduced upon quiescence or starvation (Extended data Fig. 3c and 3d), and the loss of ACF1 cannot be rescued by ATG7 deficiency (Extended data Fig. 3e), suggesting that ACF1 is not a selective autophagy substrate upon senescence. While WSTF protein is consistently lost in senescence, its mRNA levels are unchanged (Fig. 3d-3f), consistent with our proteomic and transcriptomic results (Fig. 2m and 2p). We therefore went on to investigate whether the loss of WSTF is mediated by autophagy.

**Figure 3.**
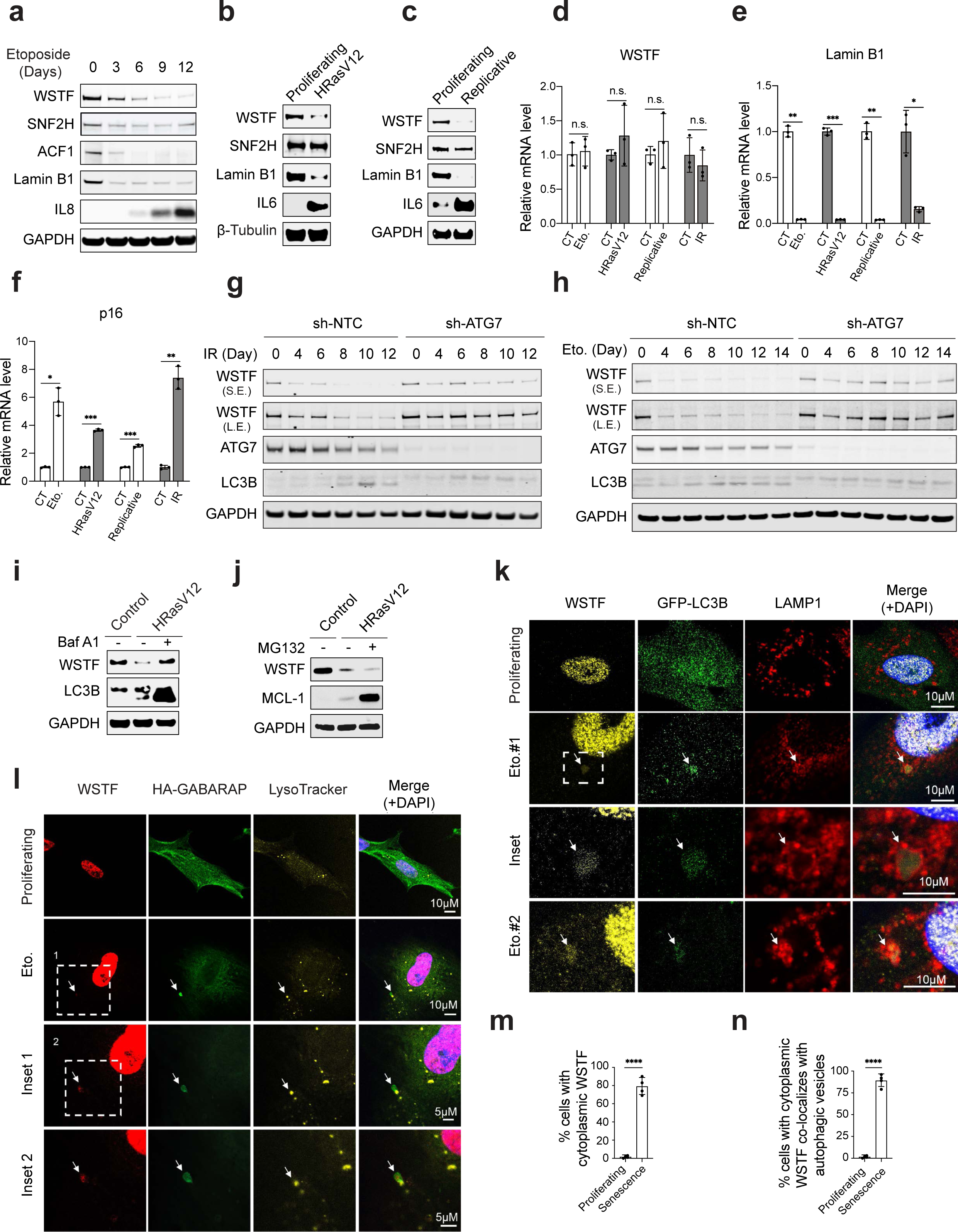
WSTF is degraded by autophagy during cellular senescence. **a**, IMR90 cells were treated with 50 µM etoposide for 2 days, then cultured in the medium without etoposide, and harvested at indicated days after the etoposide treatment. Cell lysates were subjected to immunoblotting. **b**, IMR90 cells were infected with vector control or retrovirus expressing HRasV12 for 7 days. Cell lysates were analyzed by western blotting. **c**, IMR90 cells under proliferating or replicative senescence conditions were analyzed by western blotting. **d-f**, RT-qPCR analyses of BAZ1B/WSTF (**d**), LMNB1/Lamin B1 (**e**), and CDKN2A/p16 (**f**) in proliferating, etoposide, HRasV12, replicative, or ionizing radiation (IR, 20 Gy then cultured for 14 days) induced senescence conditions. Results were normalized to Lamin A/C and presented as mean values with s.d.; n=3; p values were calculated from unpaired two-tailed Student’s t test. * P < 0.05; ** P < 0.01; *** P < 0.001; **g and h**, IMR90 cells stably expressing non-targeting control (sh-NTC) or sh-ATG7 were treated with IR (**g**) or etoposide (**h**). The cells were harvested at indicated days and analyzed by western blotting with indicated antibodies. **i and j**, IMR90 cells were infected with vector control or retrovirus expressing HRasV12 for 7 days. The cells were then treated with bafilomycin A1 (Baf A1, **i**) or MG132 (**j**) for 2 days. The lysates were analyzed by western blotting. **k**, IMR90 cells stably expressing GFP-LC3B were left untreated or induced to senesce with etoposide for a week, stained with WSTF and LAMP1 antibodies, and then imaged by confocal microscopy. WSTF signals in senescent cells were deliberately overexposed to allow our examination of its localization, and representative images are shown. Arrows indicate cytoplasmic WSTF. **l**, IMR90 cells stably expressing HA-GABARAP were incubated in growth media with 200 nM LysoTracker™ Red DND-99 for 2 h, followed by staining with WSTF and HA antibodies. The cells were imaged under a confocal microscopy. WSTF signals in senescent cells were deliberately overexposed to allow our examination of its localization, and representative images are shown. **m**, Bar graphs showing the percentage of cells with cytoplasmic WSTF in proliferating or etoposide-induced senescent cells. Data are from four randomly selected fields with over 200 cells. Results shown are mean values with s.d.; **** p < 0.0001; unpaired two-tailed Student’s t-test. **n**, Bar graph showing the percentage of cytoplasmic WSTF co-localizing with autophagy vesicles (defined as LC3B, GABARAP, LAMP1, or LysoTracker positive) in proliferating or etoposide-induced senescent cells. Results shown are mean values with s.d.; **** p < 0.0001; unpaired two-tailed Student’s t-test.

We inhibited autophagy by both genetic and pharmacological approaches, and examined the behavior of WSTF in senescence. First, we stably inactivated ATG7 using shRNA, and found that while WSTF was lost in non-targeting control shRNA cells, its protein levels were largely retained upon etoposide or IR-induced senescence in cells where ATG7 was knocked down (Fig. 3g and 3h, Extended data Fig. 3f). By contrast, the loss of ACF1 was not impaired by ATG7 deficiency (Extended data Fig. 3f). Second, we inhibited autophagy degradation using bafilomycin A1 that blocks lysosomal acidification. This led to a restoration of WSTF protein levels in senescence (Fig. 3i and Extended data Fig. 3g). By contrast, inhibition of proteasomal degradation by MG132 failed to rescue WSTF in senescence (Fig. 3j).

We further observed WSTF localization using imaging approaches. While WSTF is localized in the nucleus of proliferating control cells, cytoplasmic localization of WSTF was observed in senescence, colocalizing with autophagosome markers LC3 or GABARAP as well as lysosomal markers LAMP1 or LysoTracker (Fig. 3k and 3l, Extended data Fig. 3h and 3i, and quantified in Fig. 3m and 3n). Addition of bafilomycin A1 accumulated WSTF in the cytoplasm, colocalizing with LAMP1 (Extended data Fig. 3h). In contrast to senescence, starvation did not induce WSTF cytoplasmic localization (Extended data Fig. 3h), consistent with our immunoblotting results showing that WSTF was not lost upon starvation (Extended data Fig. 3d and 3e). Taken together, these results collectively indicate that WSTF is a selective nuclear autophagy substrate upon senescence.

### A GABARAP-WSTF interaction mediates WSTF autophagic degradation

We subsequently investigated the molecular mechanisms underlying WSTF autophagic degradation. Because ATG8 proteins bind to autophagy receptors or substrates and aid in delivering them to autophagosomes for degradation, we tested the binding between WSTF and the six ATG8 proteins. Using *in vitro* translated WSTF subjected to pull down with bacterially expressed and purified GST-ATG8 proteins, we found that GABARAP is the main ATG8 isoform that directly binds to WSTF (Fig. 4a). The binding with LC3B is quantitatively much less than that of GABARAP (Fig. 4a and 4b) in these *in vitro* pulldowns, which was further validated by co-IPs from cell extracts (Fig. 4c). These results are consistent with our ATG8 nuclear interactome data in which WSTF was detected via GABARAP co-IP (Extended data Fig. 1b and Fig. 2o). We further performed co-IP of endogenous proteins, and found that GABARAP IP brought down WSTF and its binding partner SNF2H in the ISWI chromatin remodeling complex (Fig. 4d). By contrast, GABARAP IP precipitated much less of BRG1 and BRM, components of the SWI/SNF chromatin remodeling complex^26–28^. Further examination using *in vitro* translated WSTF and SNF2H revealed that WSTF, and not SNF2H, is a direct binding partner of GABARAP (Extended data Fig. 4a and 4b). Taken together, these results strongly support a direct GABARAP-WSTF interaction.

**Figure 4.**
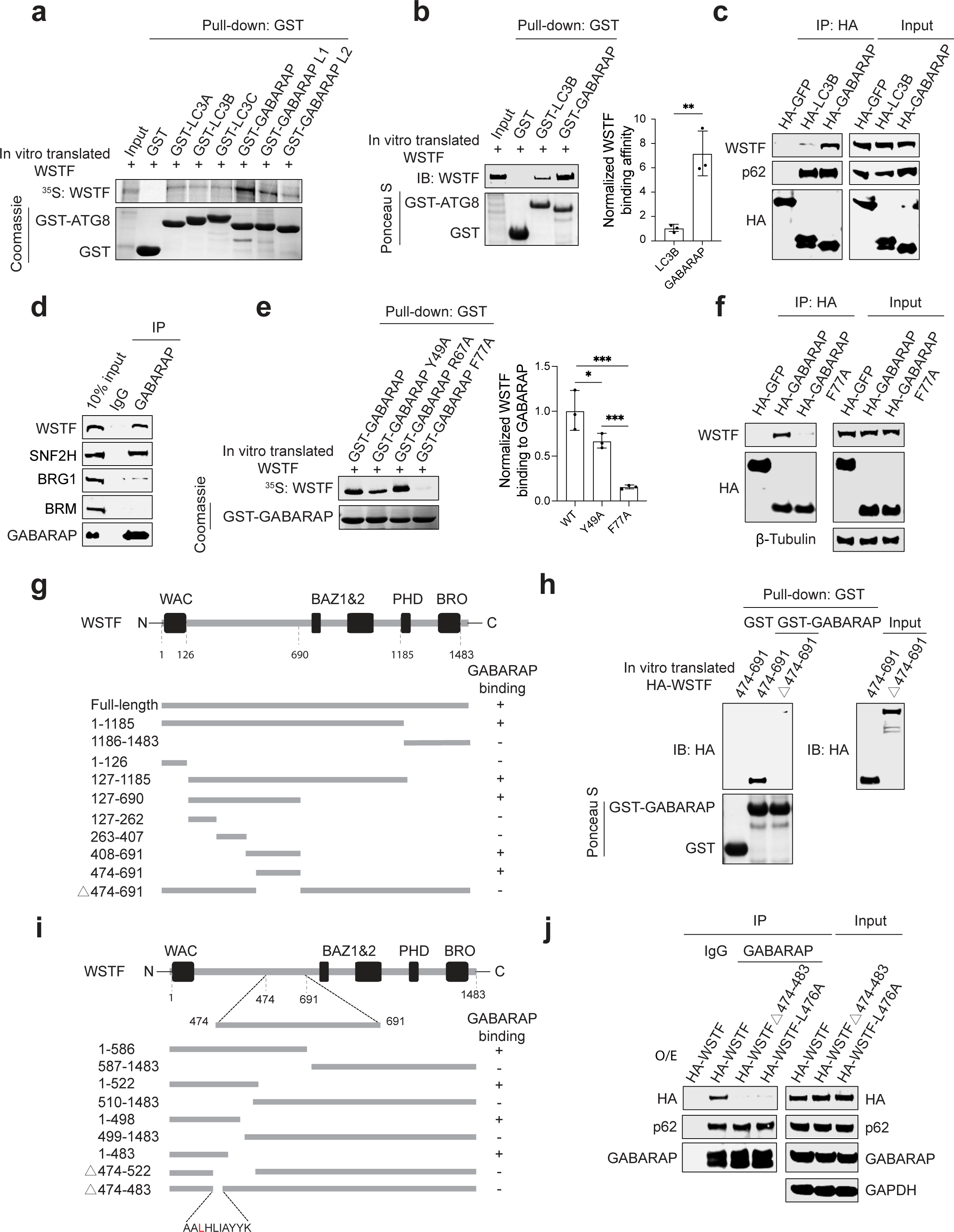
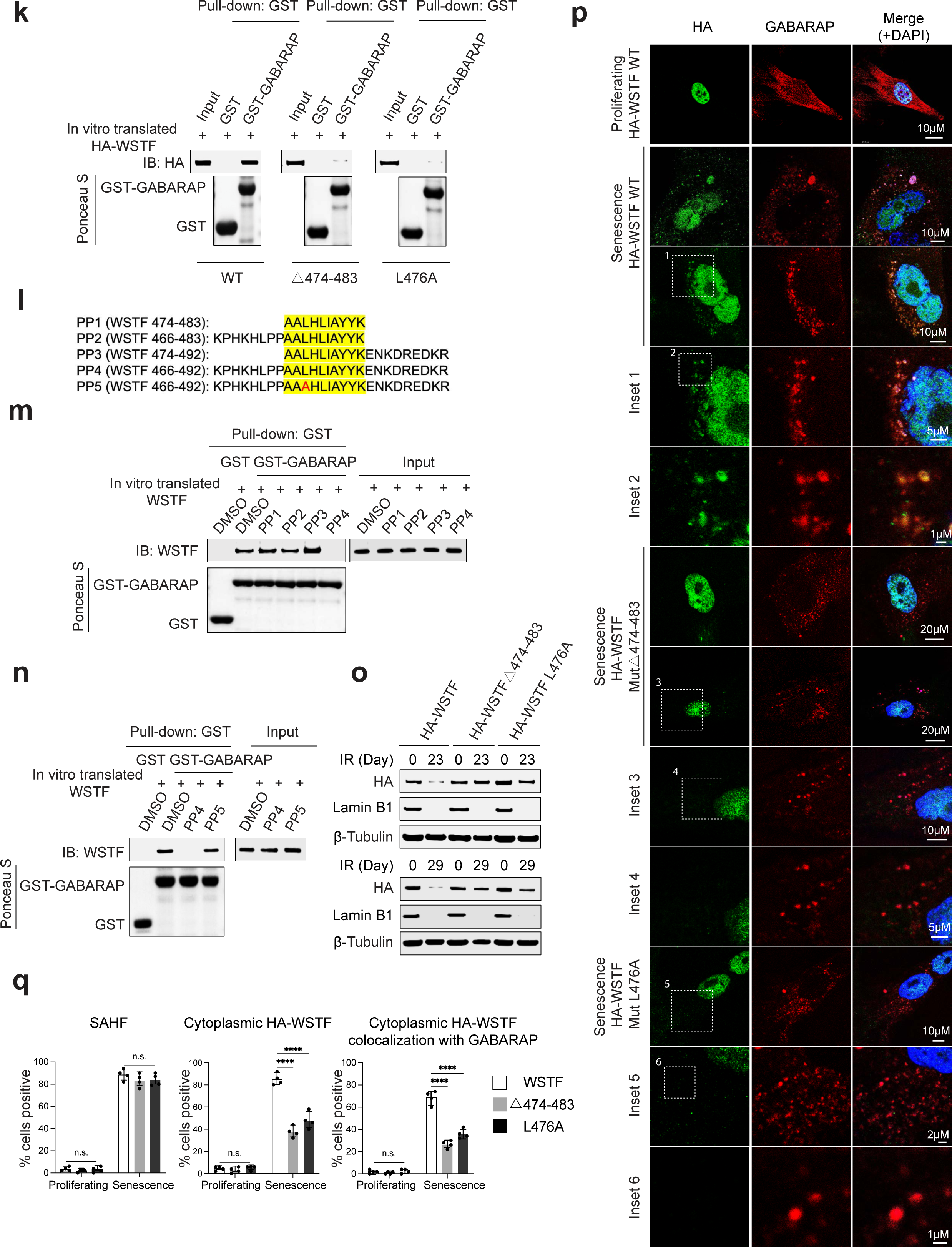
Direct binding of WSTF to GABARAP is essential for its autophagic degradation in senescence. **a**, *In vitro* translated and ^35^S-methionine labeled HA-WSTF protein was subjected to GST pull-down assays using bacterially expressed and purified GST-ATG8 proteins. **b**, *In vitro* translated WSTF protein was subjected to GST pull-down of bacterially expressed and purified GST-LC3B or GST-GABARAP. Quantification of the relative binding to WSTF is shown on the right. Data shown are mean values with s.d.; ** p<0.01; unpaired two-tailed Student’s t-test. **c**, HEK293T cells were transfected with HA-tagged constructs, followed by HA immunoprecipitation and immunoblotting with indicated antibodies. **d**, HEK293T cells were subjected to endogenous GABARAP immunoprecipitation followed by immunoblotting with indicated antibodies. **e**, *In vitro* translated and ^35^S-methionine labeled HA-WSTF protein were subjected to GST pulldown of bacterially expressed and purified GABARAP and its mutants. * P < 0.05; *** P < 0.001; one-way ANOVA coupled with Tukey’s post hoc test. **f**, HEK293T cells were transfected with HA-tagged GABARAP or its mutant and subjected to HA immunoprecipitation followed by immunoblotting with indicated antibodies. **g**, Scheme of WSTF mutants binding to GABARAP, summarizing the key findings. “+” means positive binding; “-” means negative binding. **h**, *In vitro* translated WSTF wild-type and mutants were subjected to GST-tagged GABARAP pulldown. **i,** Detailed schematic illustration of WSTF mutants in binding to GABARAP. **j**, HEK293T cells were transfected with HA-tagged WSTF constructs and subjected to GABARAP immunoprecipitation followed by immunoblotting with indicated antibodies. **k**, *In vitro* translated WSTF constructs were subjected to GST-tagged GABARAP pulldown. **l**, Scheme of synthesized peptides derived from fragments of WSTF protein, to be used in competition binding assays. **m** and **n,** *In vitro* translated WSTF protein were subjected to GST-tagged GABARAP pulldown in the presence of competing peptides illustrated in **l**. **o**, IMR90 cells were stably infected with HA-tagged WSTF constructs, subjected to IR and harvested at indicated days, then immunoblotted with indicated antibodies. **p**, IMR90 cells stably expressing HA-WSTF constructs were treated with IR and harvested at day 23, stained with HA and GABARAP antibodies, then analyzed by confocal microscopy. Representative images are shown. Note the colocalization between GABARAP and cytoplasmic WSTF in wild-type but not WSTF mutants. **q**, Related to **p**, bar graph showing the percentage of cells with senescence-associated heterochromatin foci (SAHF, measured by DAPI foci in the nucleus), cytoplasmic HA-WSTF, and cytoplasmic HA-WSTF colocalization with GABARAP. More than 400 cells in 4 fields were randomly selected and counted. Results shown are mean values with s.d.; ****P < 0.0001; one-way ANOVA coupled with Tukey’s post hoc test.

We next interrogated the regions or amino acid residues that mediate the GABARAP–WSTF interaction. On the GABARAP end, we found that the GABARAP F77A mutant significantly impaired the binding with WSTF from both *in vitro* pull-down (Fig. 4e) and co-IP from cell extracts (Fig. 4f). On the WSTF end, we constructed a series of truncations (summarized in Fig. 4g) and found that the region encompassing amino acids 474-691 of WSTF is both necessary and sufficient for interaction with GABARAP (Fig. 4h, Extended data Fig. 4c-4f). We further interrogated this region in greater detail (summarized in Fig. 4i) and found that WSTF 474-483 region, and in particular, the L476 residue, is essential for GABARAP binding in both *in vitro* pulldown and in co-IPs from cells (Fig. 4i-4k and Extended data Fig. 4g). While deletion of 474-483 or mutation of L476 abolished the binding to GABARAP, these mutants of WSTF did not affect the binding to SNF2H (mediated by the BAZ1&2 domain of WSTF) or to histone H3 (mediated by the PHD and bromodomain of WSTF) (Extended data Fig. 5a). Hence, we have discovered the critical residues on both ends that mediate the GABARAP-WSTF interaction.

**Figure 5.**
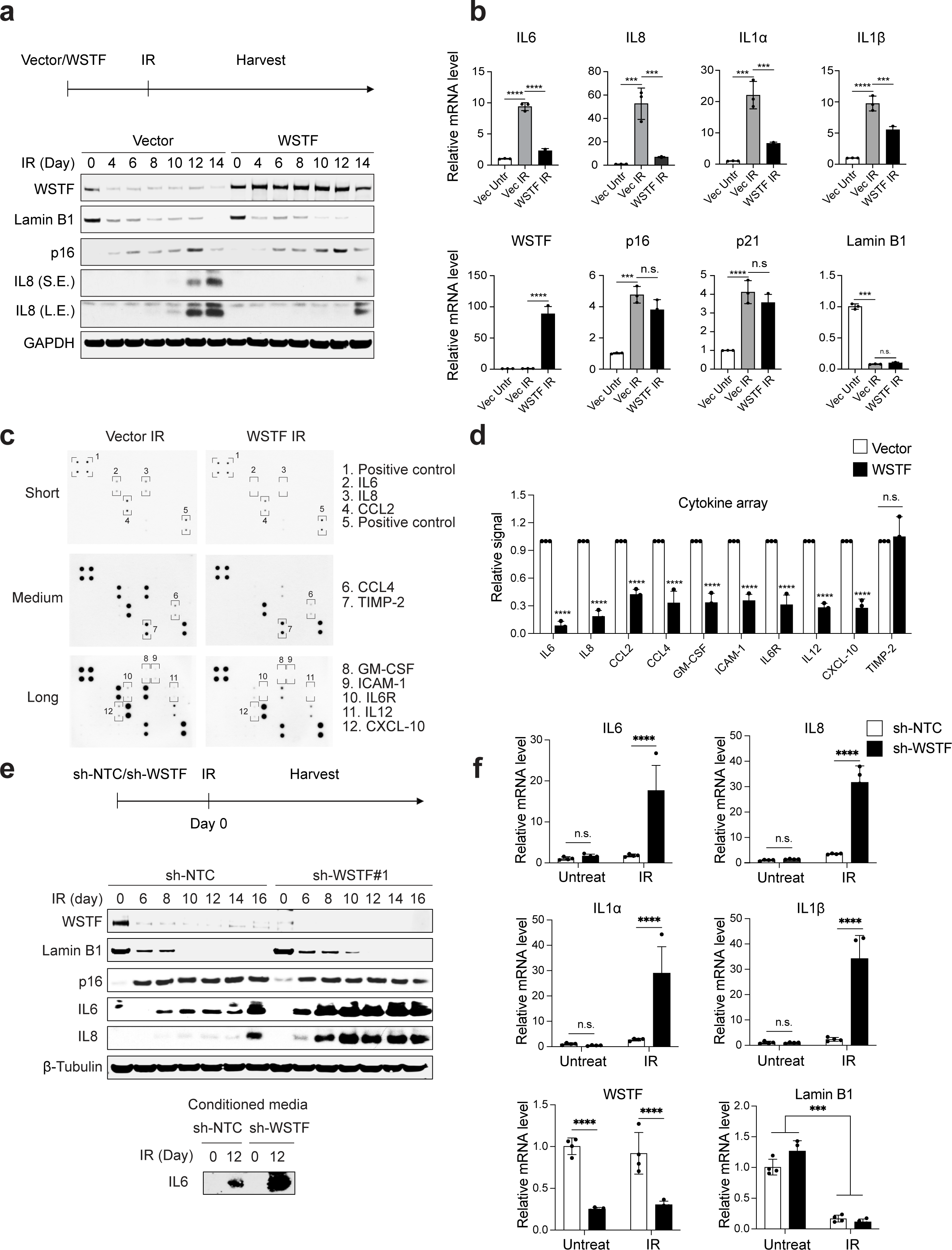
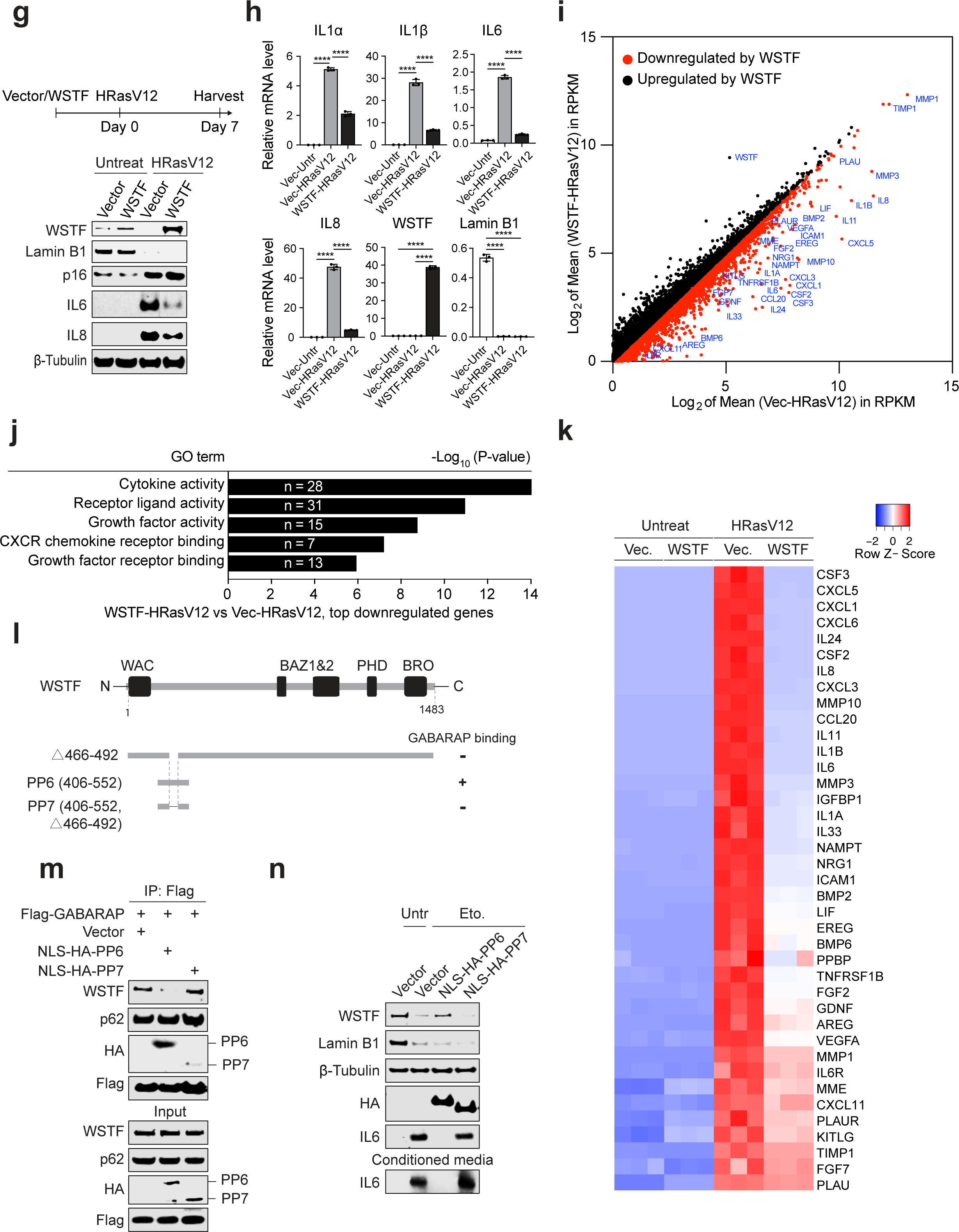
WSTF negatively regulates the SASP program of senescence. **a**, IMR90 cells stably expressing vector control or WSTF were treated by IR and harvested at indicated days, and then analyzed by western blotting. **b**, RT-qPCR analyses of IL6, IL8, ILα, IL1β, WSTF, p16, p21 and Lamin B1 in proliferating vector cells, IR vector cells and IR WSTF cells, harvested at Day 14 post IR. Results were normalized to Lamin A/C and presented as mean values with s.d.; n=3; n.s. non-significant; *** P < 0.001; **** P < 0.0001; one-way ANOVA coupled with Tukey’s post hoc test. **c**, The conditioned media from IR-induced senescence of vector or WSTF-expressing cells were analyzed by a cytokine array. Representative results with different exposures are shown. **d**, Related to **c**, quantification of the relative amounts of cytokines from the cytokine array assays. Results shown are mean ± s.e.m.; n=3; n.s. non-significant; **** P < 0.0001; unpaired two-tailed Student’s t-test. **e,** sh-NTC and sh-WSTF IMR90 cells were treated by IR and harvested at indicated days, followed by analyses of western blotting. Conditioned media were analyzed as well. **f,** RT-qPCR analyses of IL6, IL8, ILα, IL1β, WSTF, and Lamin B1 in untreated sh-NTC cells, untreated sh-WSTF cells, IR sh-NTC cells, and IR sh-WSTF cells. Cells were harvested 14 days post IR. Results were normalized to Lamin A/C and presented as mean values with s.d.; n=4; n.s. non-significant; *** P < 0.001; **** P < 0.0001; One-way ANOVA coupled with Tukey’s post hoc test. **g**, Stably overexpressed vector or WSTF IMR90 cells were infected by HRasV12 retrovirus and were harvested at indicated days. The cells were analyzed by western blotting. **h**, RT-qPCR analyses of ILα, IL1β, IL6, IL8, WSTF and Lamin B1 in untreated vector cells, HRasV12 vector cells, and HRasV12 WSTF cells. Results were normalized to Lamin A/C and presented as mean values with s.d.; n=3; **** P < 0.0001; one-way ANOVA coupled with Tukey’s post hoc test. **i,** Cells generated as in **g** were analyzed by RNA-seq. Differentially expressed genes (DEGs) were plotted in the HRasV12 condition. Representative genes up/down-regulated by WSTF in HRasV12-induced senescence are annotated. **j**, The DEGs downregulated by WSTF were analyzed by EnrichR, showing the top enriched GO biological processes with numbers of genes and P values. **k,** Heatmap presentation of the relative expression values of SASP genes. Refer to Materials and Methods for parameters used in this analysis. **l,** Scheme of WSTF peptides (PP) binding to GABARAP. “+” means positive binding; “-” means negative binding. **m,** HEK293T cells were transfected with vector, Flag-GABARAP, NLS-HA-PP6, or NLS-HAPP7 constructs and subjected to Flag immunoprecipitation followed by immunoblotting with indicated antibodies. **n,** Stably expressed vector, NLS-HA-PP6, or NLS-HA-PP7 IMR90 cells were left untreated or treated with etoposide to induce senescence and analyzed by western blotting; the conditioned media were also analyzed by western blotting.

To further investigate the interaction, we synthesized a series of WSTF peptides corresponding to the WSTF 474-483 region (Fig. 4l), and asked whether these peptides could block the GABARAP-WSTF interaction. The peptide corresponding to 474-483 of WSTF along with its flanking sequence at the amino and carboxy (N and C) termini (PP4, residues 466-492) effectively inhibited GABARAP-WSTF binding in *in vitro* pulldown (Fig. 4m). Introducing L476A mutation into this peptide giving PP5 abrogated this effect (Fig. 4n).

Several interaction surfaces are known for binding to ATG8 family proteins with the LC3-interacting region (LIR) docking site (LDS) being the most used surface for interaction partners^29^. Recently, another binding surface named ubiquitin-interacting motif (UIM)-docking site (UDS) was suggested^30^. Using point mutations, we can compromise the two binding surfaces selectively (Extended data Fig. 5b). We used the Y49A mutation to render the LDS binding deficient and the F77A mutation to cripple the UDS in GABARAP. Our results clearly implicate the UDS in binding to WSTF (Fig. 4e and 4f). We next investigated the structure of the WSTF region binding to GABARAP using *in silico* modeling. The 474-484 region of WSTF is predicted to form an alpha-helix, similar to a helix of the CaV2.2 protein^31^, based on PHYRE2 server that predicts protein structure^32^ (Extended data Fig. 5c). Furthermore, structural modeling of WSTF by AlphaFold2 ^33, 34^ predicts with high confidence that the 473-485 region of WSTF forms an alpha-helix, with the L476 residue exposed at the outer surface of WSTF (Extended data Fig. 5d). Employing the ClusPro web server for automatic protein-protein docking^35^, we obtained a top ranked model placing the WSTF helix with the L476 residue firmly into the UDS of GABARAP (Extended data Fig. 5d). The UDS surfaces of LC3B and GABARAP are different with a wider and more shallow pocket in GABARAP than in LC3B, which helps explain why WSTF binds GABARAP with higher affinity than LC3B (Extended data Fig. 5e, Fig. 4b). Together, these results provided mechanistic insights into the WSTF-GABARAP interaction and prompted us to investigate the behavior of WSTF mutants in senescence.

We stably expressed the WSTF △474-483 mutant or the WSTF L476A mutant defective in binding to GABARAP, and investigated their degradation and localization. While the protein levels of wild-type WSTF were reduced upon senescence, both mutants showed impaired downregulation in senescence, measured at two time points post IR (Fig. 4o). By contrast, the loss of Lamin B1 was not affected by the two mutants (Fig. 4o). In addition, we found that while wild-type WSTF was translocated to the cytoplasm, colocalizing with GABARAP puncta, both mutants showed impaired ability to do so (Fig. 4p and quantified in Fig. 4q, Extended data Fig. 5b). The WSTF mutants did not affect the formation of senescence-associated heterochromatin foci (SAHF), a marker for senescence (Fig. 4q). Taken together, these results strongly suggest that WSTF binding to GABARAP is required for its autophagic degradation in senescence.

### WSTF represses the SASP program

We subsequently investigated the biological significance of autophagic degradation of WSTF in senescence. To counteract the loss of WSTF, we overexpressed WSTF, and examined the key features of senescence (Fig. 5a). We found that WSTF overexpression did not affect the loss of Lamin B1 (Fig. 5a), induction of p16Ink4a (hereafter referred to as p16) (Fig. 5a), or senescence-associated beta-galactosidase (SA-β-gal) (Extended data Fig. 6a). However, the induction of IL8, a SASP factor, was reduced by WSTF overexpression (Fig. 5a). RT-qPCR analyses of senescence and SASP genes confirmed that while WSTF overexpression did not affect Lamin B1, p16, or p21, the induction of key factors of the SASP program was significantly reduced, including IL6, IL8, IL1α, and IL1β (Fig. 5b). The impaired SASP program was further confirmed at the secreted level, measuring the conditioned media of senescent cells overlaid to a cytokine array (Fig. 5c). While multiple SASP factors showed reduced levels in the conditioned media, TIMP-2, a soluble factor constitutively secreted by proliferating and senescent IMR90 cells, was not altered by WSTF overexpression (Fig. 5c and quantified in Fig. 5d). In addition to WSTF overexpression, we also asked whether disruption of WSTF promotes the SASP program. shRNA-mediated WSTF inactivation, using two independent hairpins, enhanced SASP gene expression, while not affecting Lamin B1 or p16 upon IR-induced senescence (Fig. 5e and 5f, Extended data Fig. 6b and 6c). Furthermore, we found that WSTF overexpression blocks the SASP program of oncogene-induced senescence triggered by HRasV12 (Fig. 5g and 5h). Moreover, expression of WSTF in established senescent cells that already lost WSTF inhibited IL8 expression (Extended data Fig. 6d). Taken together, these data collectively suggest that WSTF is a negative regulator of the SASP program.

**Figure 6.**
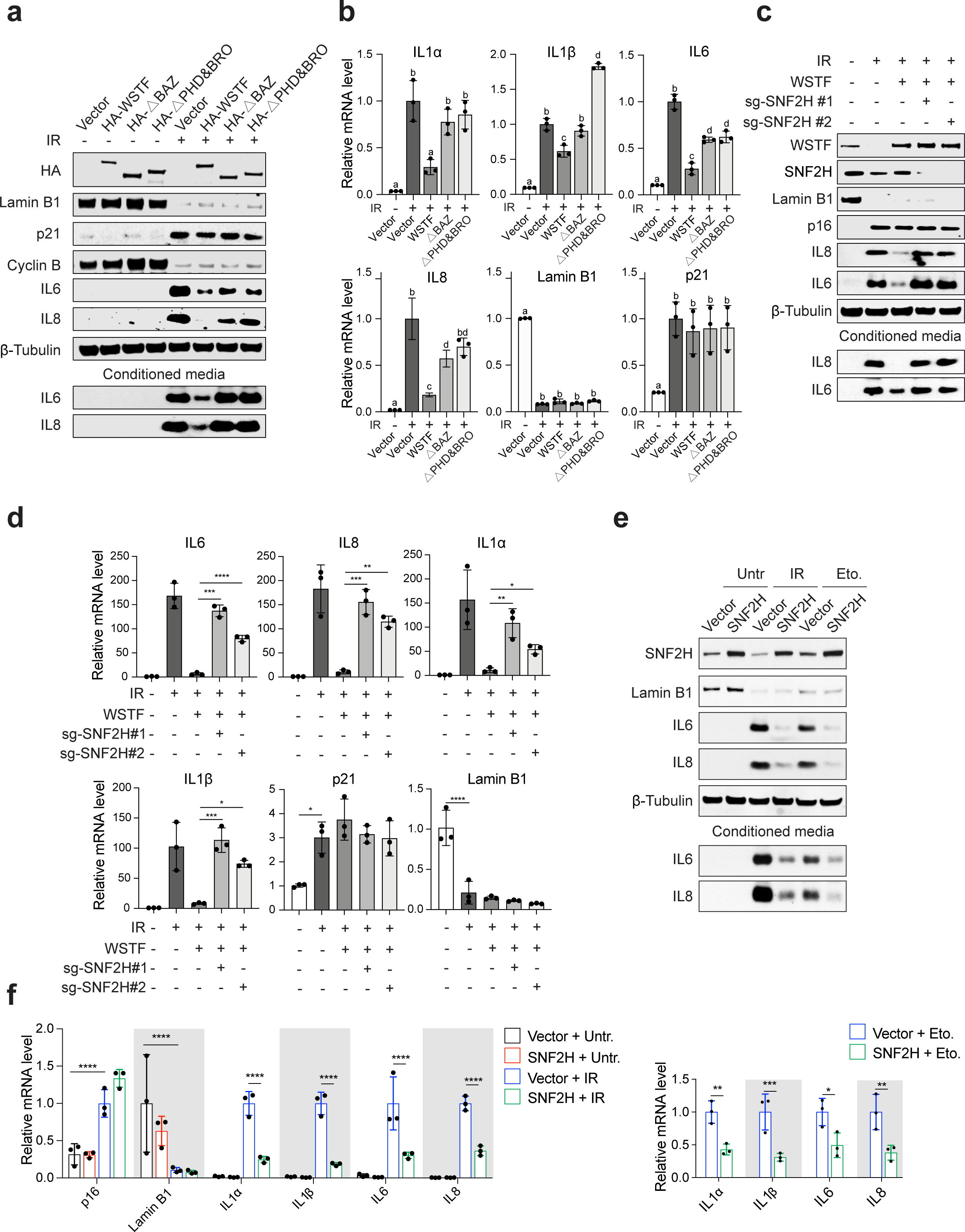
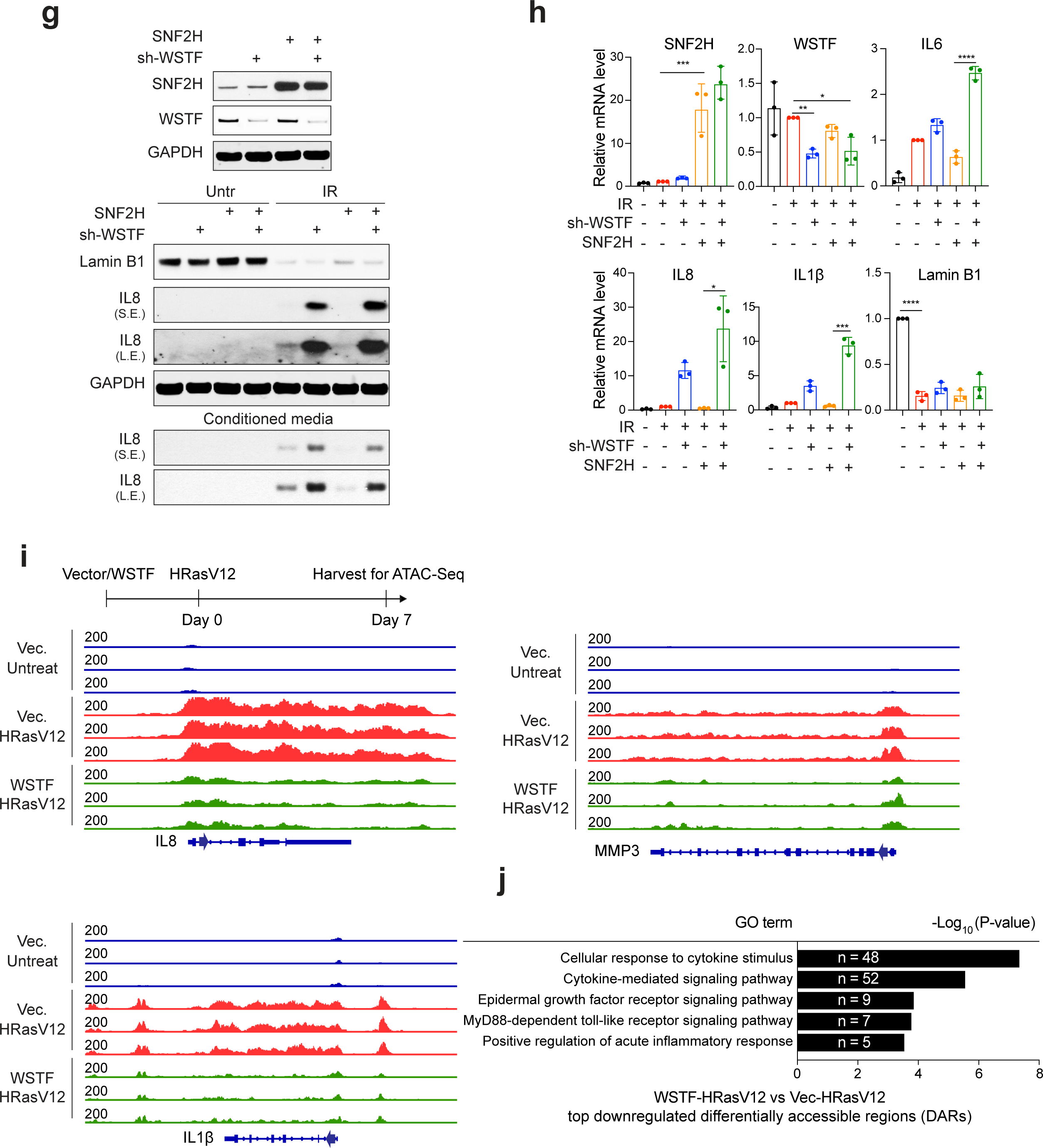
WSTF inhibits chromatin accessibility over SASP genes. **a**, Full-length WSTF or WSTF truncations were stably expressed in IMR90 cells and were left untreated or induced to senescence by IR. Cells were harvested at day 14 post IR and analyzed by western blotting (**a**) or RT-qPCR (**b**). **b**, RT-qPCR analyses of ILα, IL1β, IL6, IL8, Lamin B1 and p21. Results were normalized to Lamin A/C and presented as mean values with s.d.; n=3; letters (e.g., a, b, and c) to highlight significant differences. Groups that are not significantly different are assigned a common letter. In other words, two treatments without a common letter are statistically significant at the chosen level of significance (P ≤ 0.05), one-way ANOVA coupled with Tukey’s post hoc test. **c and d,** SNF2H inactivation by CRISPR abrogates the effect of WSTF overexpression. IMR90 cells were engineered to express control sg-RNA or sg-RNA against SNF2H, combined with vector or WSTF overexpression. The cells were left untreated or induced to senescence by IR and were harvested at Day 14. The samples were analyzed by immunoblotting (**c**) or RT-qPCR (**d**). **d**, RT-qPCR analyses of IL6, IL8, ILα, IL1β, p21 and Lamin B1. Results were normalized to Lamin A/C and presented as mean values with s.d.; n=3; * P < 0.05; ** P < 0.01; *** P < 0.001; one-way ANOVA coupled with Tukey’s post hoc test. **e and f**, Vector and SNF2H-overexpressing cells were left untreated or induced to senescence by IR or etoposide. The samples were analyzed by western blotting (**e**) or RT-qPCR (**f**). **f**, RT-qPCR analyses of SNF2H, p16, Lamin B1, ILα, IL1β, IL6, and IL8. Results were normalized to Lamin A/C and presented as mean values with s.d.; n=3; ** P < 0.01; *** P < 0.001; **** P < 0.0001; one-way ANOVA coupled with Tukey’s post hoc test. **g and h,** IMR90 cells were engineered to overexpress SNF2H in combination with WSTF knockdown, and were left untreated or induced to senescence by IR. The samples were analyzed by immunoblotting (**g**) or RT-qPCR (**h**). **h**, RT-qPCR analyses of IL6, IL8, IL1β, SNF2H, WSTF and Lamin B1. Results were normalized to Lamin A/C and presented as mean values with s.d.; n=3; n.s. non-significant; * P < 0.05; ** P < 0.01; *** P < 0.001; **** P < 0.0001; one-way ANOVA coupled with Tukey’s post hoc test. **i and j,** ATAC-seq analyses of vector control and WSTF-overexpressing cells. **i,** Scheme of experimental design is noted on top, and ATAC-seq genomic tracks of IL8, IL1β and MMP3 are presented. **j**, Enrichment analysis showing the top GO categories enriched among genes at or proximal to differential accessible regions (DARs) whose chromatin accessibility decreased in WSTF-HRasV12 compared to Vec-HRasV12. The numbers of genes corresponding to the DARs and the P values are shown.

To further examine the SASP program in an unbiased manner, we performed an RNA-seq study, including proliferating control and HRasV12-induced senescence, expressing vector control or WSTF (following the design of Fig. 5g). Analysis of differentially expressed genes (DEGs) in senescent cells suggested that WSTF substantially inhibited the expression of a large panel of pro-inflammatory genes (Fig. 5i). Pathway enrichment analysis using EnrichR^36^ revealed that genes downregulated by WSTF are strongly enriched in the categories of cytokines, growth factors, and chemokines, characteristic of the SASP program (Fig. 5j and specific genes listed in Extended data Fig. 6e). Genes related to other processes like cell cycle arrest are not affected by WSTF. The global reduction of the SASP gene expression is also presented in a heatmap, including established SASP factors^37^ (Fig. 5k). These results strongly support that WSTF specifically represses the SASP program of senescence.

Lastly, we asked whether inhibition of WSTF autophagic degradation affects the SASP. To this end, we expressed a short region of WSTF (406-552) that binds to GABARAP, using deletion of the core amino acids for GABARAP interaction 466-492 as a negative control (illustrated in Fig. 5l). We tagged these peptides with a nuclear localization signal (NLS), leading to their expression in the nucleus (Extended data Fig. 6f). While WSTF 406-552 peptide strongly inhibited the binding between full-length WSTF and GABARAP, likely due to its competition with WSTF for GABARAP binding, WSTF 406-552 △466-492 showed impaired binding to GABARAP, accompanied by its inability to block full-length WSTF-GABARAP interaction. Upon induction of senescence, the 466-492 peptide inhibited WSTF downregulation and the expression of IL6, while the 406-552 △466-492 mutant peptide failed to do so. Hence, these data indicate that autophagic degradation of WSTF, mediated by GABARAP, is critical for the induction of the SASP program of senescence.

### WSTF suppresses chromatin accessibility of SASP genes

We next investigated how WSTF suppresses the SASP program of senescence. We first asked whether WSTF affects the established mechanisms of the SASP^21, 38–45^, and found that while WSTF blocked the SASP gene expression (Extended data Fig 7a), it did not alter the status of p38MAPK, DNA damage response, mTOR, cytoplasmic chromatin fragments (CCF), cGAS-STING pathway, NF-κB p65 subunit nuclear translocation and phosphorylation at S536, or senescence-associated heterochromatin foci (SAHF) (Extended data Fig. 7a to 7c). These data suggest that WSTF employs a previously uncharacterized mechanism to regulate the SASP program.

**Figure 7.**
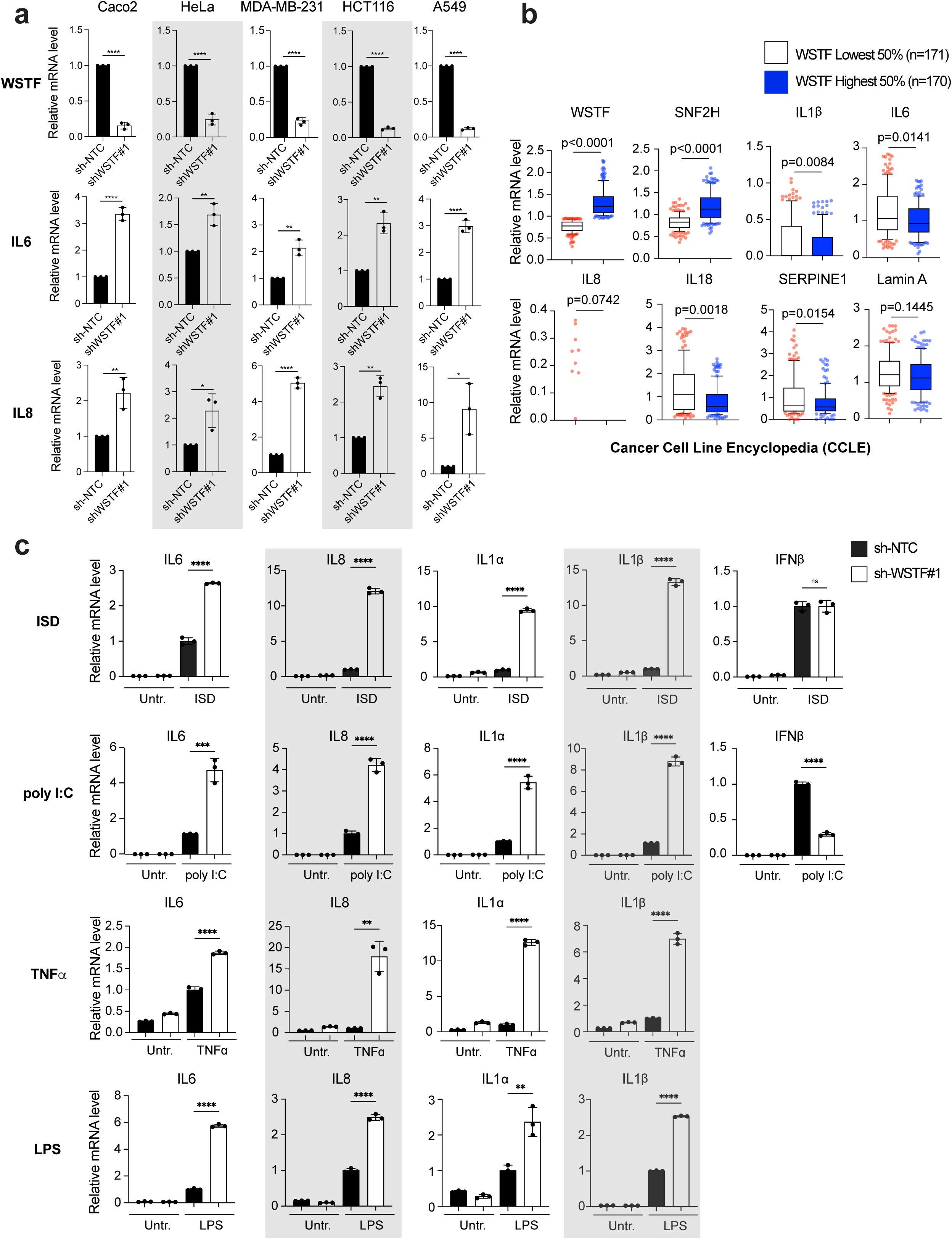
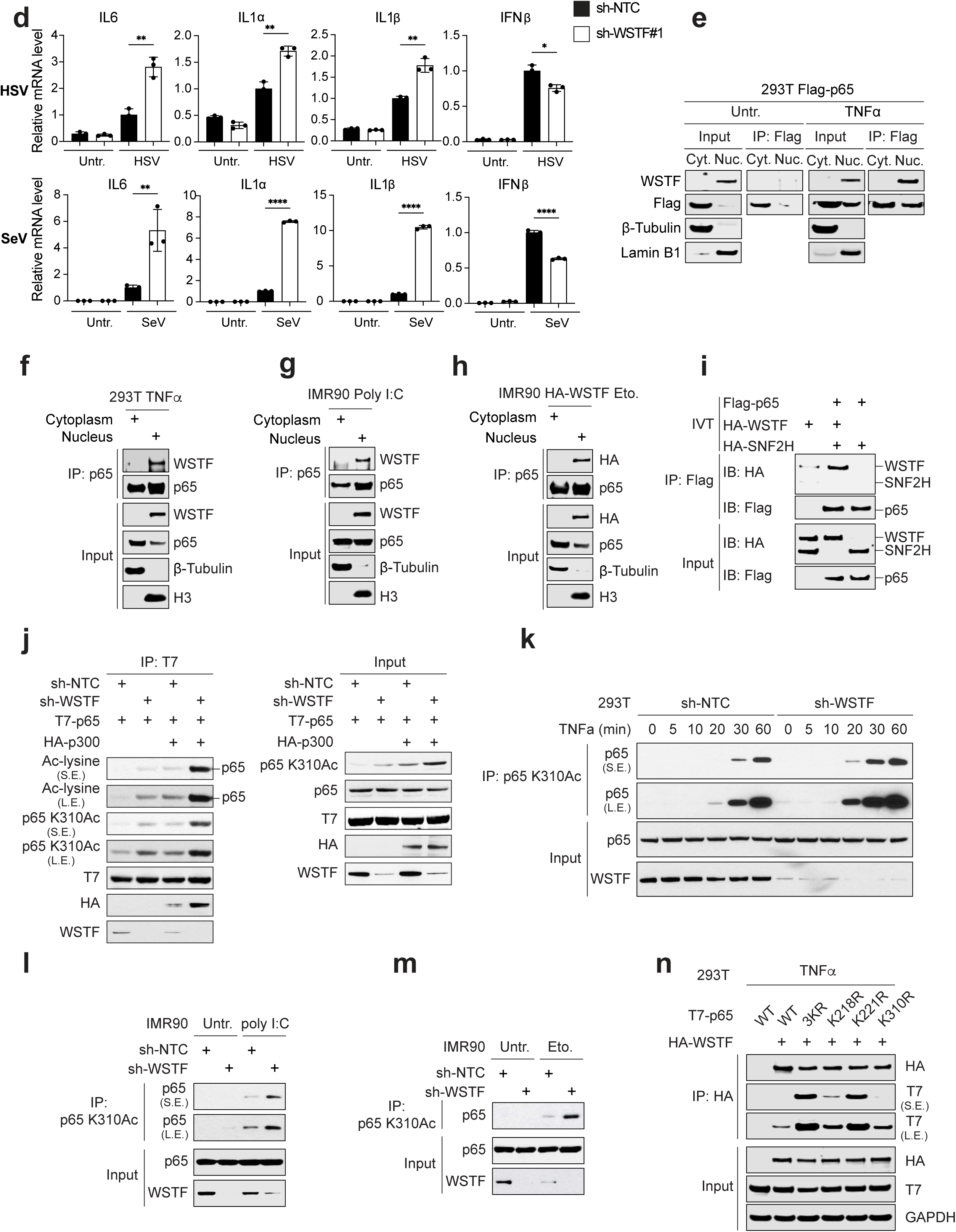
WSTF is a broad inhibitor of NF-κB-mediated pro-inflammatory gene expression. **a**, RT-qPCR analyses of IL6, IL8, and WSTF in sh-NTC and sh-WSTF stably expressing cancer cells, including CaO.2, HeLa, MDA-MB-231, HCT116, and A549. Results were normalized to Lamin A/C and presented as mean values with s.d.; n=3; n.s. non-significant; * P < 0.05; ** P < 0.01; *** P < 0.001; **** P < 0.0001; unpaired two-tailed Student’s t-test. **b**, Cancer Cell Line Encyclopedia (CCLE) analyses of WSTF and inflammatory gene correlation, which contains proteomic datasets from over 300 cancer cell lines. Based on the protein levels of WSTF, we grouped these cancer cell lines into the lowest 50% subset and the highest 50% subset, then compared the expression of SNF2H, IL1β, IL6, IL8, IL18, SERPINE1 and Lamin A between these two subsets. For box plots displayed in this study, the central rectangle spans a range from the first quartile to the third quartile, also called the Interquartile Range. A line inside the rectangle shows the median. The whiskers are drawn down to the 10th percentile and up to the 90th. Points below and above the whiskers are drawn as individual dots. Outliers were defined as data points that were either 1.5 × interquar-tile range or more above the third quartile, or 1.5 × interquartile range or more below the first quartile. Unpaired t test with Welch’s correction (do not assume equal SDs) was used to compute statistical significance. P values were shown as indicated. **c**, RT-qPCR analyses of IL6, IL8, ILα, IL1β, and IFNβ in sh-NTC and sh-WSTF BJ-hTERT cells for ISD, poly I:C, TNFα stimulation, and THP1 cells for LPS stimulation. Results were normalized to Lamin A/C and presented as mean values with s.d.; n=3; n.s. non-significant; * P < 0.05; ** P < 0.01; *** P < 0.001; **** P < 0.0001; unpaired two-tailed Student’s t-test. **d,** RT-qPCR analyses of IL6, ILα, IL1β and IFNβ in sh-NTC and sh-WSTF BJ-hTERT cells for SeV infection, and THP1 cells for HSV infection. Results were normalized to Lamin A/C and presented as mean values with s.d.; n=3; n.s. non-significant; * P < 0.05; ** P < 0.01; *** P < 0.001; **** P < 0.0001; unpaired two-tailed Student’s t-test. **e,** HEK293T cells were transfected with a Flag-p65 construct and treated with or without TNFα (20 ng/ml) for 30min, then fractionated into cytoplasm and nucleus, followed by Flag immunoprecipitation and immunoblotting with indicated antibodies. **f,** HEK293T cells were treated with TNFα and fractionated into cytoplasm and nucleus, then subjected to p65 immunoprecipitation at the endogenous level, followed by immunoblotting with indicated antibodies. **g,** IMR90 cells were treated with poly I:C (30 ug/ml) for 1.5h, and fractionated into cytoplasm and nucleus, then subjected to p65 immunoprecipitation at the endogenous level, followed by immunoblotting with indicated antibodies. **h,** IMR90 cells stably expressing HA-WSTF were treated with etoposide to induce senescence, and fractionated into cytoplasm and nucleus, then subjected to p65 immunoprecipitation and immunoblotting with indicated antibodies. **i,** *In vitro* translated Flag-p65, HA-WSTF, and HA-SNF2H were subjected to Flag immunoprecipitation and immunoblotting with indicated antibodies. Note that WSTF binds quantitatively more to p65 than that of SNF2H. **j,** HEK293T cells stably expressing sh-NTC or sh-WSTF were transfected with constructs encoding T7-p65 or HA-p300, then subjected to T7 immunoprecipitation and immunoblotting with indicated antibodies. **k,** HEK293T cells stably expressing sh-NTC or sh-WSTF were treated with TNFα for indicated minutes, then subjected to p65 K310Ac immunoprecipitation and immunoblotting with indicated antibodies. **l and m,** IMR90 cells stably expressing sh-NTC or sh-WSTF were transfected with poly I:C (**l**) or induced to senescence by etoposide (**m**). The cells were subjected to p65 K310Ac immunoprecipitation and immunoblotting with indicated antibodies. **n,** HEK293T cells were transfected with constructs encoding HA-WSTF, T7-p65 wild-type or its mutants (K218R, K221R, K310R, and 3KR), treated with TNFα, and then subjected to HA immunoprecipitation and immunoblotting with indicated antibodies.

We truncated the domains of WSTF with established biological functions and asked which activities of WSTF are involved in the SASP program. WSTF harbors a WAC domain with tyrosine kinase activity^46^, a BAZ1&2 domain that binds to SNF2H, and PHD & bromodomains that bind to histones (illustrated in Extended data Fig. 7d). We generated a series of truncation mutants of WSTF, and confirmed that the BAZ1&2 truncation impaired the binding to SNF2H while the PHD & bromodomain truncation impaired the binding to histones (Extended data Fig. 7d and 7e). Upon induction of senescence, we found that the BAZ1&2 domain as well as PHD & bromodomains are required for WSTF to inhibit the SASP (Fig. 6a and 6b). By contrast, the WAC domain of WSTF is not required to inhibit the SASP (Extended data Fig. 7f and 7g). These results suggest that the binding of SNF2H and association with chromatin are necessary for WSTF to suppress the SASP program of senescence.

We thus investigated the connection with SNF2H in greater detail, and discovered that SNF2H-WSTF complex is critical in repressing the SASP. First, CRISPR-mediated inactivation of SNF2H abrogated the effect of WSTF in blocking the SASP (Fig. 6c and 6d), consistent with the results that the BAZ1&2 domain of WSTF is involved (Fig. 6a and 6b). Second, overexpression of SNF2H inhibited the SASP (Fig. 6e and 6f), suggesting that SNF2H, like WSTF, is also a repressor of the SASP. Third, inactivation of WSTF abrogated the effect of SNF2H in blocking the SASP (Fig. 6g and 6h). Together, these results indicate that the ISWI chromatin remodeling complex has a previously unappreciated role in suppressing the SASP program of senescence.

Because the ISWI complex represses gene expression by forming ordered nucleosomes on chromatin, we assessed whether WSTF regulates chromatin accessibility over SASP genes by ATAC-seq^47^. Chromatin accessibility at SASP genes was low in control cells but was massively increased in senescence (Fig. 6i showing genomic tracks of ATAC-seq read density at IL8, MMP3, and IL1β). Importantly, expression of WSTF substantially reduced chromatin accessibility at these SASP genes (Fig. 6i). We subsequently performed peak calling and evaluated the differentially accessible regions (DARs) at the genome-wide scale. GO analysis of the DARs revealed strong enrichment of functional gene categories involving cytokines, growth factors, and acute inflammation, characteristics of the SASP program (Fig. 6j and specific genes listed in Extended data Fig. 8a). The global reduction of chromatin accessibility at representative SASP genes is also presented in a heatmap (Extended data Fig. 8b). Further, the DARs with reduced chromatin accessibility located at the genes with downregulated RNA-seq expression in senescence were strongly enriched with genes of the SASP program (Extended data Fig. 8c). In sum, our results strongly suggest that WSTF in cooperation with SNF2H suppresses chromatin accessibility of pro-inflammatory genes of senescence.

**Figure 8.**
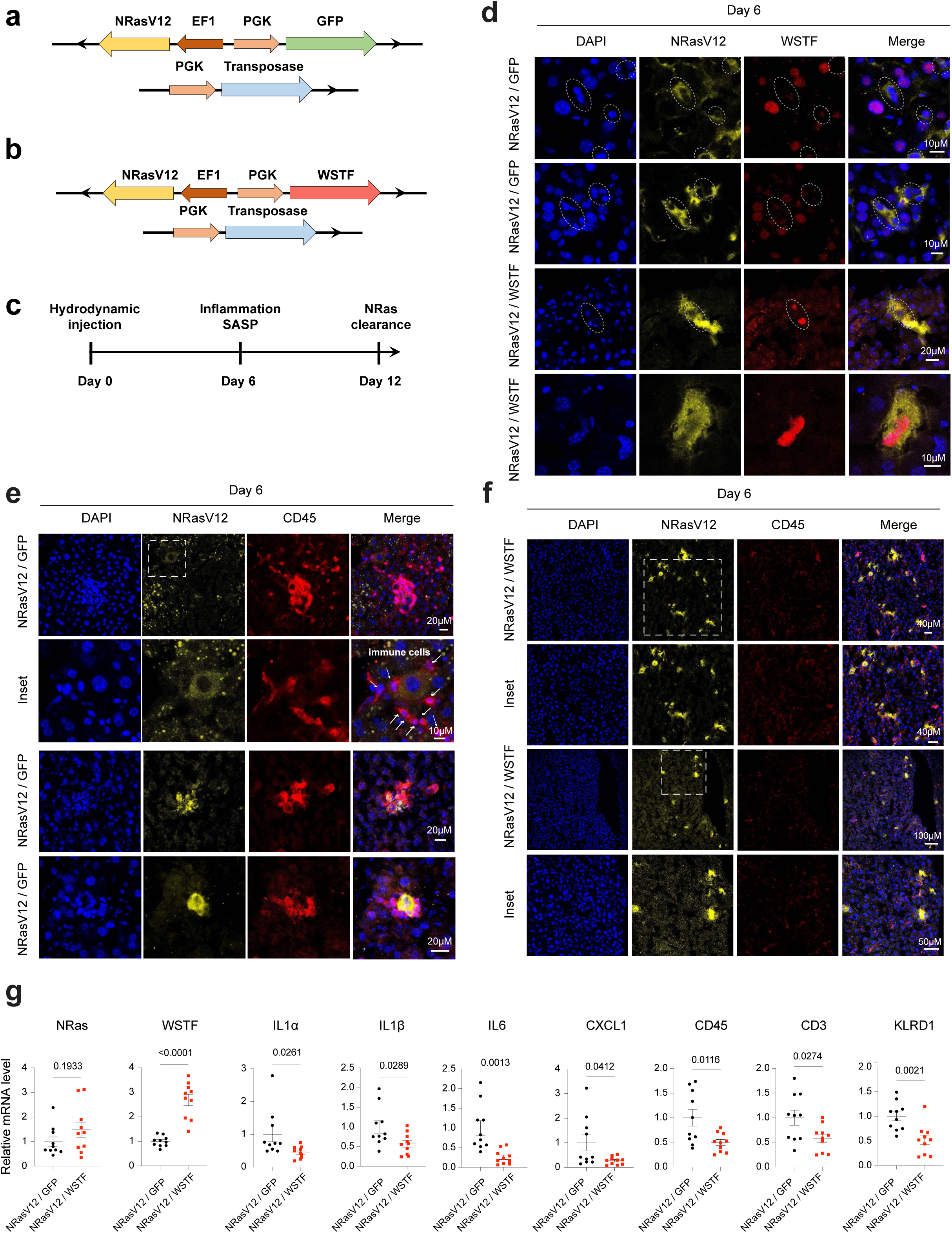
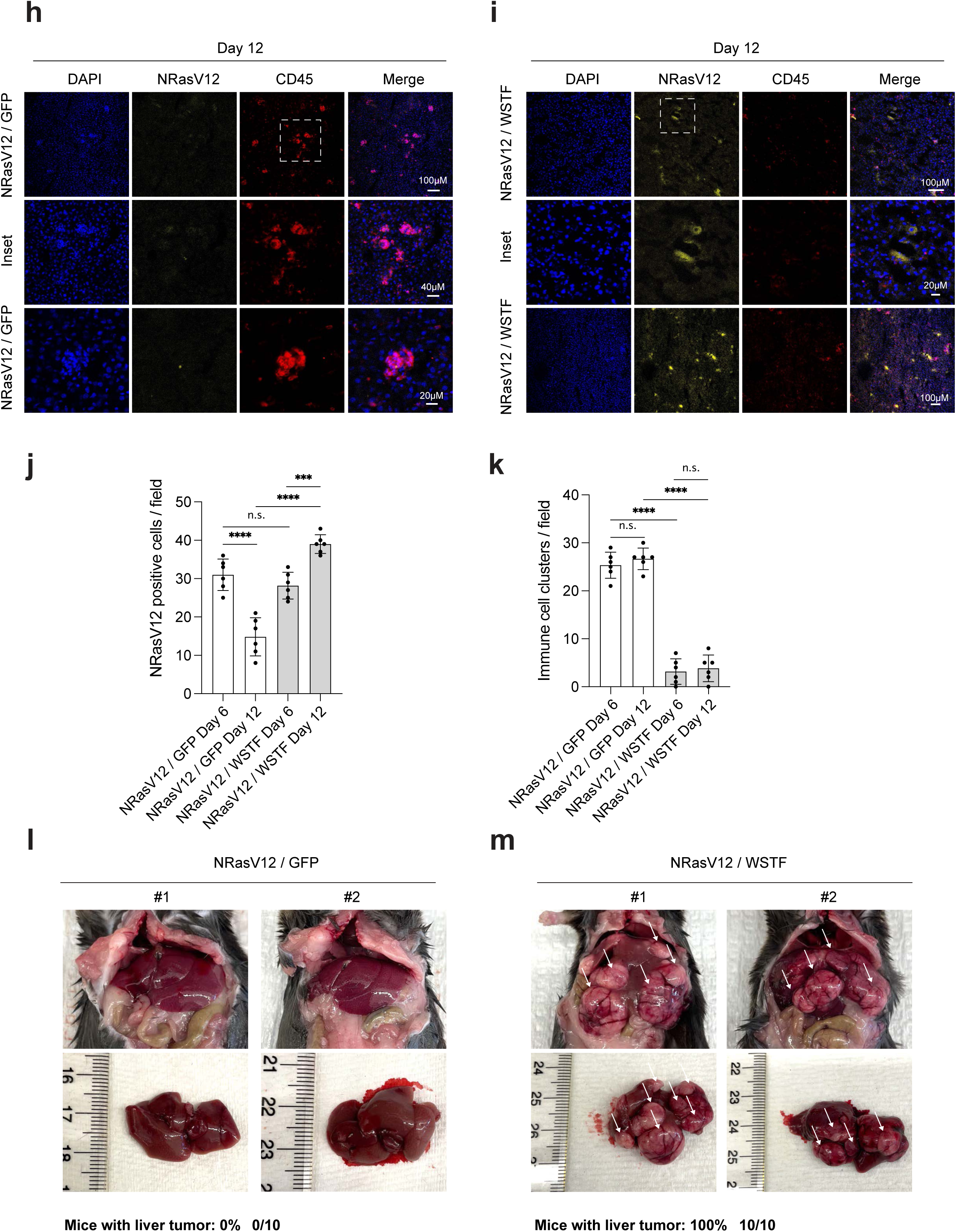
Loss of WSTF is required for immunosurveillance of activated Ras in mouse liver. **a**, **b**, **c**, Schematic illustration of constructs and experimental design. **d**, Liver sections 6 days post injection were stained with NRasV12 and WSTF antibodies, followed by imaging under a confocal microscopy. NRasV12-positive cells were highlighted. Note the expression level of WSTF was reduced in the GFP group and increased in the WSTF group. **e** and **f**, Liver sections 6 days post injection were stained with NRasV12 and CD45 antibodies, then analyzed by a confocal microscopy. The immune cells are highlighted with arrows. Scale bars are shown as indicated. **g**, RT-qPCR analyses of liver from the GFP and WSTF groups. The relative expression levels of pro-inflammatory genes and immune cell genes were measured. 10 mice in each group were used, P values were shown as indicated, calculated by unpaired two-tailed Student’s t-test. **h** and **i,** Liver sections 12 days post injection were stained with NRasV12 and CD45 antibodies, then analyzed by a confocal microscopy. Scale bars were shown as indicated. **j**, Bar graphs showing the quantification of NRasV12 positive cells in different experimental groups and presented as mean values with s.d.; n = 6; n.s.: non-significant. *** P < 0.001; **** P < 0.0001; one-way ANOVA coupled with Tukey’s post hoc test. **k**, Bar graph showing the quantification of immune cell clusters in different experimental groups and presented as mean values with s.d.; n = 6; n.s.: non-significant. **** P < 0.0001; one-way ANOVA coupled with Tukey’s post hoc test. **l** and **m,** Representative images of liver tumors from NRasV12/GFP and NRasV12/WSTF groups, 6 months after the injection. The numbers of mice with liver tumors are shown at the bottom.

### WSTF is a broad inhibitor of inflammation

Senescence shares several features with cancer. Senescent cells induced by activated oncogenes or loss of tumor suppressor proteins can generate neoplastic lesions *in vivo*. Both senescence and cancer are associated with DNA damage, and they share similar features of genome methylation^48^. Using cancer cell line encyclopedia (CCLE), we previously showed that the cGAS-STING pathway, required for the SASP program of senescence, is also strongly linked to cancer-associated pro-inflammatory gene expression^21^. These connections prompted us to investigate whether WSTF is involved in pro-inflammatory gene expression in cancer.

We inactivated WSTF using shRNA, and found that loss of WSTF increased the expression of IL6 and IL8 in Caco2, HeLa, MDA-MB-231, HCT116, and A549 cells (Fig. 7a and Extended data Fig. 9a). To robustly examine the connection between WSTF and pro-inflammatory gene expression in cancer at a larger scale, we exploited the proteomic resource of CCLE database that contains the proteomes of over 300 cancer cell lines^49^. Cell lines with lowest and highest 50% of WSTF expression were grouped and the expression of key pro-inflammatory genes was compared between WSTF-low and WSTF-high subsets. This analysis revealed that higher WSTF expression is linked to higher expression of SNF2H, and importantly, lower expression of pro-inflammatory genes, while the housekeeping gene Lamin A does not follow this pattern (Fig. 7b). Together with our WSTF knockdown in cancer cell results, these data collectively suggest that WSTF is a negative regulator of pro-inflammatory genes in broad scenarios of cancer cells.

**Figure 9.**
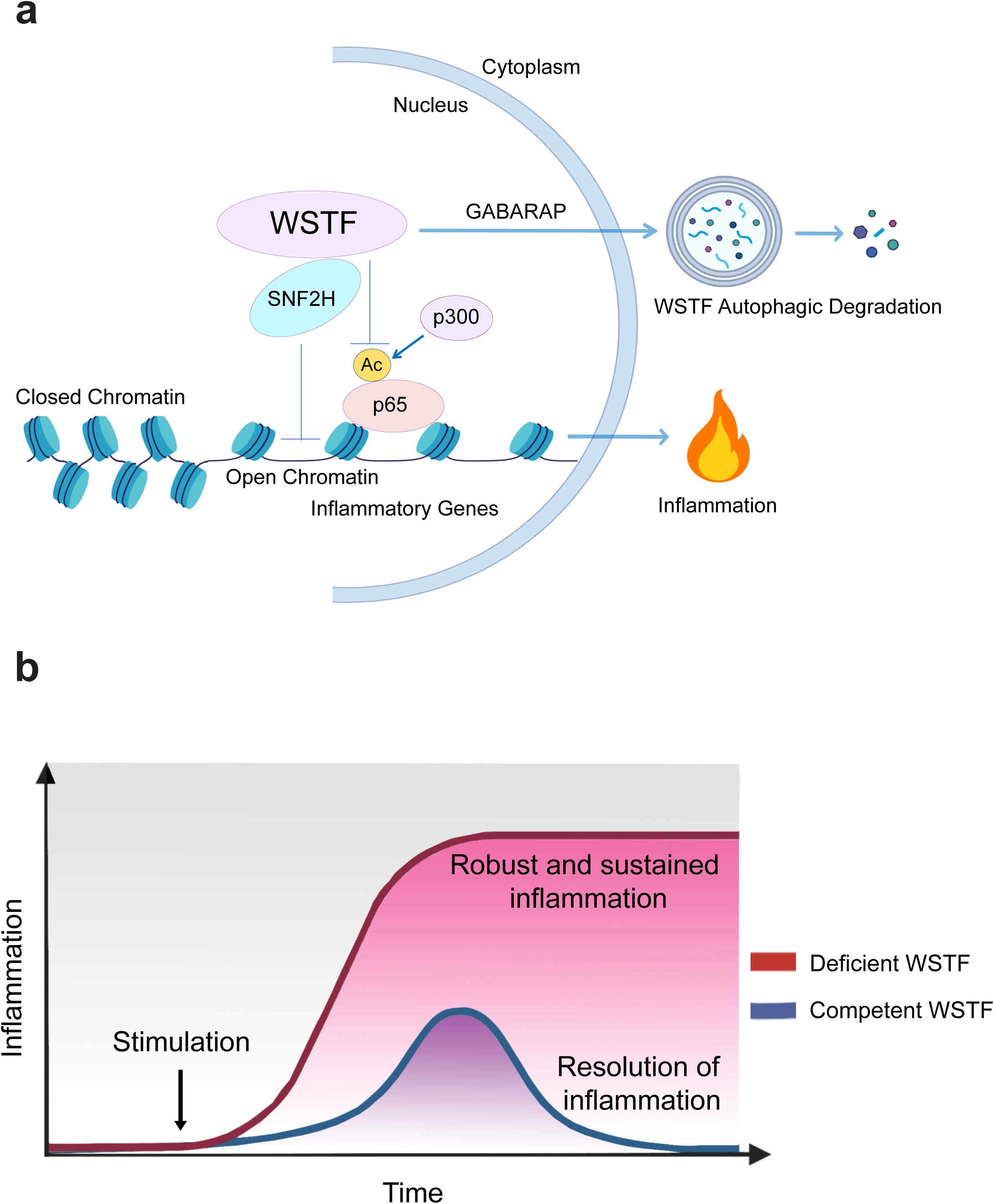
Schematic illustration for the roles of WSTF in regulating inflammation. **a**, Autophagic degradation of WSTF induces inflammation. WSTF, via SNF2H, represses chromatin accessibility over inflammatory genes. In addition, WSTF competes with p300 for p65 binding, inhibiting p65 acetylation and transcription of inflammatory genes. WSTF binding to GABARAP leads to nucleus-to-cytoplasm translocation of WSTF and its autophagic degradation, which promotes inflammation. **b,** Biological roles for WSTF in regulating inflammation. In WSTF competent conditions, inflammation can be induced and properly resolved in a timely manner. In WSTF-deficient conditions, rapid and robust inflammation is induced, which causes a longer period of sustained inflammation.

The results in cancer prompted us to examine a hypothesis that WSTF is a broad inhibitor of inflammatory gene expression. We exposed the control and WSTF deficient cells with a variety of pro-inflammatory conditions, including dsDNA (ISD) and dsRNA (poly I:C) transfection, tumor necrosis factor alpha (TNFα), and bacterial lipopolysaccharides (LPS). In all these conditions, WSTF deficiency led to significantly higher expression of pro-inflammatory cytokines (Fig. 7c and Extended data Fig. 9b-9d). Moreover, we examined viral infection, using DNA virus (HSV-1) and RNA virus (SeV). In both cases, WSTF deficiency led to significantly higher expression of pro-inflammatory genes (Fig. 7d). The induction of interferon gene (IFNβ) was not upregulated by WSTF deficiency (Fig. 7c and 7d). Together with our results in senescence and cancer, these data strongly suggest that WSTF is a broad negative regulator of inflammation.

### WSTF directly binds to p65 and inhibits p65 acetylation

The specific inhibition of inflammatory genes by WSTF, in a variety of contexts, sparked us to further examine the molecular mechanisms by which WSTF blocks inflammation. While we presented results showing that WSTF inhibits chromatin accessibility of SASP genes in senescence, it remains unclear what instructs WSTF to specifically affect inflammatory genes. A clue is provided from our RNA-seq and ATAC-seq data, exploring the transcription factors associated with the downregulated genes by WSTF. In both datasets, a strong enrichment of NF-κB and RelA (p65) subunit of NF-κB was detected (Extended data Fig. 10a and 10b). We subsequently investigated how WSTF is connected to NF-κB, a key transcription factor for inflammatory genes.

We first asked whether WSTF interacts with the p65 subunit of NF-κB. In basal condition, p65 is in the cytoplasm while WSTF is in the nucleus, thus an interaction was not detected; upon stimulation by TNFα, p65 is translocated to the nucleus, and an interaction with WSTF was detected by co-IP (Fig. 7e). The WSTF-p65 interaction was further confirmed by co-IPs at the endogenous level upon TNFα (Fig. 7f) or poly I:C stimulation (Fig. 7g), and in WSTF overexpressed senescent cells (Fig. 7h, of note, endogenous WSTF is lost in senescent cells and thus overexpressed WSTF was used). Importantly, using *in vitro* translated WSTF and p65, a direct interaction was detected (Fig. 7i). By contrast, p65 failed to bind to *in vitro* translated SNF2H (Fig. 7i). In sum, we found that WSTF of the ISWI complex directly binds to the p65 subunit of NF-κB.

We went on further to examine whether WSTF regulates p65 activity. In basal condition, p65 binds to IκB in the cytoplasm. Upon inflammatory stimulation, IκB is phosphorylated, leading to its proteasomal degradation and p65 translocation to the nucleus^50^. Nuclear p65 undergoes a series of modifications, such as phosphorylation and acetylation, facilitating p65 binding to chromatin and stimulation of the transcription of inflammatory genes^51^. We found that p65 nuclear translocation and phosphorylation at S536 were not affected by WSTF deficiency (Extended data Fig. 10c-10g), consistent with the behavior of p65 in senescent cells overexpressed with WSTF (Extended data Fig. 7a and 7b).

An important modification of p65 is the reversible acetylation^52^. Acetylation of p65, catalyzed by p300 or CBP, stimulates p65 binding to DNA and its transcription activity^53^. Acetylation at K310 of p65 is important for recruitment of BRD4 and transcription of inflammatory genes ^54, 55^. Using T7-tagged p65 coupled with WSTF knockdown and p300 overexpression, we found that WSTF deficiency substantially increased p65 overall acetylation and acetylation at K310 (referred to as K310ac, Fig. 7j). Consistent with the increased acetylation of p65, enhanced p65 binding to p300 was detected (Fig. 7j). We further examined p65 K310ac at the endogenous level, and found that WSTF deficiency increased K310ac upon TNFα stimulation (Fig. 7k), accompanied by increased mRNA levels of IL6 and IL8 (Extended data Fig. 10h). Likewise, K310ac was increased in WSTF-deficient cells upon poly I:C stimulation (Fig. 7l) and in senescence (Fig. 7m).

Lastly, we examined the binding between WSTF and acetylated p65. In addition to K310, other important sites of acetylation have also been reported, including K218 and K221^53^. Mutation of the three acetylation sites to R (termed as 3KR)^53^ substantially increased the binding to WSTF upon TNFα stimulation (Fig. 7n). Using individual K to R mutations, we found that K221 is the predominant residue that mediates the interaction with WSTF (Fig. 7n). These data suggest that WSTF preferentially binds to unmodified lysine residues on p65 to prevent p65 acetylation. We thus envision two-step events for NF-κB activation in the nucleus (illustrated in Extended data Fig. 10i). Upon p65 translocation to the nucleus, WSTF directly binds to non-acetylated p65 and inhibits p300 binding, serving as an inhibitor of p65 acetylation and activation (Extended data Fig. 10i, left). When stimulation further propagates, p300 eventually acetylates p65, leading to WSTF dissociation from p65 and transcription activation by NF-κB (Extended data Fig. 10i, right).

### Loss of WSTF is required for immuno-surveillance of activated Ras *in vivo*

We investigated the importance of WSTF in regulating inflammation *in vivo*. An important role of inflammation is to restrain tumorigenesis in response to oncogene activation. The SASP program triggered by activated oncogenes can alarm the immune system to induce immuno-surveillance and clearance of the pre-malignant cells, acting as a potent tumor suppressive mechanism^11, 12^. In particular, many primary cell types cope with oncogenic Ras activation by inducing cellular senescence, which, via the SASP, activates an initial state of tumor suppressive inflammation^11, 12^. An established model for oncogene-induced senescence and the SASP in mice is liver expression of NRasV12, which induces hepatocyte senescence, the SASP, immune cell infiltration, and clearance of the NRasV12-expressing hepatocytes^56^. We previously reported that ablation of STING blocked the SASP program, leading to impaired immuno-surveillance and formation of liver tumors^21^.

A vector with a Sleeping Beauty transposon that co-expresses NRasV12 and GFP, together with a transposase construct, was co-injected into the mouse tail vein through hydrodynamic injection (illustrated in Fig. 8a). WSTF was cloned to co-express with NRasV12 to investigate the role of WSTF in this system (Fig. 8b). This approach leads to specific and stable hepatocyte expression of NRasV12 together with GFP or WSTF (Fig. 8a and 8b). The liver was analyzed 6 days after injection for inflammation and the SASP, and 12 days after injection for NRasV12 hepatocyte clearance (outlined in Fig. 8c).

NRasV12 expression in the liver led to specific loss of WSTF in NRasV12-positive hepatocytes, but not in NRasV12-negative cells (Fig. 8d and Extended data Fig. 11a), consistent with our results that oncogene-induced senescence lost WSTF in primary human cells (Fig. 3b). We therefore overexpressed WSTF together with NRasV12 in the same vector (Fig. 8b and 8d) to investigate the SASP and inflammation in the liver.

6 days post injection, NRasV12/GFP mice induced infiltration of CD45-positive immune cells surrounding NRasV12-positive hepatocytes (Fig. 8e). Clusters of immune cells can also be observed by DAPI staining (Fig. 8e). These responses are consistent with what we and others previously reported in this model^21, 56^. By sharp contrast, NRasV12/WSTF mice showed dramatically impaired infiltration of immune cells and formation of immune clusters, although NRasV12 was expressed at comparable levels to the NRasV12/GFP group (Fig. 8f and quantified in Fig. 8j and 8k). In addition, we performed RT-qPCR analyses to quantify the transcripts of SASP genes and immune cell genes. The critical pro-inflammatory factors IL1α, IL1β, IL6, and CXCL1 were all significantly reduced in WSTF group, accompanied by reduced CD45 (a marker for all immune cells), CD3 (T cells), and KLRD1 (NK cells) (Fig. 8g). These results indicate that NRasV12-induced loss of WSTF is required for the induction of SASP and inflammation in the liver.

12 days post the injection, NRasV12-positive hepatocytes showed reduced positivity, due to immune-mediated clearance of the pre-malignant cells (Fig. 8h and quantified in Fig. 8j and 8k). By contrast, NRasV12/WSTF group exhibited persistence of NRasV12-positive hepatocytes, accompanied by reduced infiltration of CD45-positive immune cells (Fig. 8i and quantified in Fig. 8j and 8k). Impaired clearance of NRasV12-positive hepatocytes can lead to malignant growth of these cells and progression to liver tumors that express NRasV12^21, 56^. Six months following the hydrodynamic injection, while none of the GFP group developed liver tumors (0 out of 10) (Fig. 8l and Extended data Fig. 11b), severe intrahepatic tumors were observed in all mice of the WSTF group (10 out of 10) (Fig. 8m and Extended data Fig. 11c). These data strongly indicate that loss of WSTF is a critical event to induce immuno-surveillance of oncogenic Ras in mice.

## Discussion

In this study, we generated a nuclear ATG8 interaction network in primary human fibroblasts and in mouse brain, uncovering hundreds of novel interactions. Combined with the nuclear proteomes in senescence, we identified WSTF as a previously unknown nuclear substrate of autophagy. Autophagic degradation of WSTF, via interaction with GABARAP, stimulates senescence-associated inflammation, by promoting chromatin accessibility over inflammatory genes. In addition to senescence, we uncovered that WSTF is a broad repressor of inflammation, via direct binding to the NF-κB p65 subunit and inhibiting its acetylation. Lastly, we demonstrated that loss of WSTF is essential for immuno-surveillance of activated oncogene in mice. A schematic model summarizing our findings is presented in Fig. 9a.

Our study offers new perspectives and insights in several fields of research. First, our nuclear ATG8 interactome offers a systematic resource for studying nuclear autophagy. While ATG8 interactomics studies have been conducted before^15–17^, we reason that, due to technical issues in lysing nuclear and chromatin fractions (Fig. 1a), prior resources were in fact limited to cytoplasmic interactions. Our nuclear autophagy interaction network differs substantially from a previously established autophagy-interaction network^15^ (Extended data Fig. 1c), which we hope will spark other investigators to explore nuclear autophagy, an area still in its infancy. Second, our proteomic datasets in senescence provide a new resource to investigate senescence. We present a whole cell proteome in HRasV12-induced senescence as well as nuclear proteomes of etoposide-induced senescence, in wild-type and ATG7-deficient cells. Together with our RNA-seq datasets in prior^7, 21^ and current studies, these resources will allow investigators to discover new RNA and protein alterations in senescence. Third, we uncovered a new mechanism regulating senescence-associated inflammation (the SASP program). While the roles of SASP in cancer, wound healing, and aging have been documented^11, 12^, the mechanisms that initiate the robust and sustained inflammation are not fully understood. We showed that WSTF altering chromatin accessibility over inflammatory genes is a new mechanism to initiate the SASP program. This mechanism is distinct from cGAS-STING and other established mechanisms of the SASP. Fourth, we unveiled a novel activity of the ISWI chromatin remodeling complex. While WSTF and ISWI complex have been reported to be involved in regulating development and other biological processes^57–60^, our study is the first to connect ISWI complex with senescence and inflammation, thus suggesting a new strategy to target inflammatory disorders. Lastly and importantly, this study discovers a novel regulatory mechanism of NF-κB and inflammation, from the nucleus (described below).

Inflammation is considered a “double-edged sword”. Although inflammation is essential for restraining infection and tumorigenesis, chronic and uncontrolled inflammation contributes to autoimmunity and many other diseases, including most age-associated diseases^61^. Hence, investigation of negative regulators of inflammation is of critical importance. Our discovery of WSTF as a negative regulator of NF-κB sheds a new light on the complex role of inflammation. We showed that while WSTF competent cells are able to properly resolve inflammation, WSTF deficient cells, such as in senescent cells, exhibited robust and sustained inflammation (illustrated in Fig. 9b). Mechanistically, besides the ISWI activity, WSTF binds to p65 in the nucleus, suppressing its acetylation and activation. This role of WSTF is distinct from other reported negative regulators of p65 in the nucleus, such as enzymes that remove acetylation or phosphorylation marks of p65^51^, which require p65 to be activated first. By contrast, WSTF preferentially binds to unmodified lysine residues of p65 (Fig. 7n), preventing p65 activation. This mechanism of WSTF is similar to IκB in the cytoplasm, serving as a constitutive nuclear inhibitor of NF-κB. Hence, WSTF acts as a “checkpoint” to raise the threshold of signals that can produce inflammatory gene expression. This ensures that (1) only bona-fide inflammatory signals can produce successful inflammation, and (2) inflammation can be properly resolved to prevent tissue damage. We noted that the pro-inflammatory conditions under acute setting did not show loss of WSTF, unlike the loss of WSTF in cellular senescence which takes days to weeks to establish. Future studies are needed to investigate autophagic degradation of WSTF under chronic inflammatory conditions besides senescence. We envision that WSTF is likely to be involved in other inflammatory disorders, such as age-associated diseases, metabolic disorders, and cytokine-associated toxicity. Our study further suggests that targeting WSTF autophagic degradation may hold promise to intervene in inflammation-associated diseases.

## Acknowledgements

The nuclear autophagy interactome studies in human fibroblasts were conducted in collaboration with J. Wade Harper and Joseph D. Mancias. We thank Maria Grazia Vizioli for contributing part of the western blots in Fig. 3a as well as Yuting Tan for generating the sgRNA constructs targeting SNF2H and the next-gen sequencing samples in control cells. We thank Yanxin Xu, Takuya Kumazawa, Yaosi Liang, and Yuting Tan for technical support and discussions. We acknowledge the microscopy core facility of Center for Regenerative Medicine at Massachusetts General Hospital for assistance on confocal microscopy, and the next-gen sequencing core at Massachusetts General Hospital for assistance on RNA-seq. We thank Riley Graham for advice on ATAC-seq. Z.D. is supported by National Institute of Health (NIH) awards R35GM137889, R00AG053406, UG3CA268117, R21AG073894, and Glenn Foundation for Medical Research and AFAR Grant for Junior Faculty. T.J. is supported by a strategic thematic grant from UiT and by TOPPFORSK (grant 249884) program of the Research Council of Norway. Work by J.L. and S.H.S. is supported in part by the Harvard Stem Cell Institute and Harvard Catalyst NIH UL1TR002541.

## Author contributions

Z.D. conceived the project. Y.W. conducted most of the experiments unless stated. V.V.E. contributed the mass-spectrometry results of ATG8 nuclear interactomes and proteomes in human fibroblasts, performed in J. Wade Harper and Joseph D. Mancias laboratories. A.K. contributed part of the GABARAP-WSTF interaction results. A.O. contributed part of the autophagy and senescence data. X.L., X.Z., and Z.Y. contributed mouse brain LC3 interactome. M.C. and R.I.S. contributed computational analyses of RNA-seq and ATAC-seq. L.W. contributed experimental design and part of the next-gen sequencing data. J.L. and S.H.S contributed part of the protein interaction maps. C.B. and P.D.A. contributed part of the mouse liver tissue samples and *in vivo* data. Z.D. generated samples for next-gen sequencing in senescent cells and contributed part of the autophagy, senescence, and inflammation results. Z.D. and Z.Z. conceived and supervised inflammation experiments. S.V.S. and R.E.K. contributed novel reagents of ISWI complex and supervised epigenetic experiments. T.J. and Z.D. supervised the study and provided funding support. Y.W., T.J. and Z.D. wrote the paper. All authors discussed the manuscript.

The authors declare no competing financial interests.

## Materials and methods

### Cell culture and treatment

Primary IMR90, BJ fibroblasts and MEFs were described previously^6, 21^. The cells were cultured in DMEM supplemented with 10% fetal bovine serum (FBS), 100 units/mL penicillin, and 100 μg/mL streptomycin (Invitrogen), and were intermittently cultured with plasmocin (Invivogen) and tested for mycoplasma using MycoAlert^TM^ PLUS Mycoplasma Detection Kit (Lonza). The cells were cultured under physiological oxygen (3%), and were used within population doubling of 35, except for replicative senescence experiments where cells were cultured till replication exhaustion (around population doubling 80). For etoposide-induced senescence, IMR90 cells at approximately 60–70% confluency were treated with 50 uM etoposide for 48 hours and harvested at days indicated in figure legends. To establish senescence of BJ cells, the cells at approximately 60-70% confluency were treated with 40 uM etoposide for 24 h and harvested at days indicated in figure legends. For ionizing radiation (IR), cells were irradiated with X-Rad 320 (Precision X-Ray Irradiator) at 20 Gy and harvested as described in figure legends. For HRasV12-induced senescence, retrovirus from retroviral vector encoding HRasV12 was used to infect cells. Following hygromycin selection, cells were cultured for 7 or more days as described in figure legends. These conditions reproducibly induced senescence, confirmed by cell cycle arrest measured by lack of Cyclin A and phosphorylated Rb, EdU less than 5% positivity, and SA-β-gal over 95% positivity, as described in our previous studies (Dou et al., 2017; Dou et al., 2015; Vizioli et al., 2020; Xu et al., 2020). HeLa (American Type Culture Collection; CCL2) and A549 (ATCC CCL185) were grown in DMEM (Sigma-Aldrich; D5796) supplemented with 10% FBS (Biochrom AG; S0615) and 1% streptomycin-penicillin (Sigma-Aldrich; P4333). HEK293T and MEFs were grown in DMEM (Sigma-Aldrich; D6046) supplemented with 10% FBS and 1% streptomycin-penicillin. Caco2 cells (ATCC HTB-37) from a colorectal adenocarcinoma were grown in DMEM (Sigma-Aldrich; D6046) supplemented with 10% FBS and 1% streptomycin-penicillin. HCT116 cells (ATCC CCL-247) from a colorectal carcinoma were grown in McCoy’s 5A (Sigma-Aldrich; M9309) supplemented with 10% FBS and 1% streptomycin-penicillin. MDA-MB-231 cells (ATCC HTB-26) from a breast adenocarcinoma were grown in high glucose DMEM (Sigma-Aldrich; D6046) supplemented with 10% FBS, 2mM glutamine and 1% streptomycin-penicillin. 0.2 µM BafA1 or 10 µM MG132 were used in experiments described in Fig. 3i and 3j.

### Mice experiments

All mice experiments were performed in compliance with the Institutional Animal Care and Use Committee at Icahn School of Medicine at Mount Sinai and Massachusetts General Hospital. GFP-LC3 transgenic mice were described elsewhere^18^. Both sexes were included for mice studies. Hydrodynamic tail vein injection was performed as previously described^21^. In brief, 20 μg of NRasV12 and 10 μg of transposase constructs were injected to C57BL/6 mice that were 8– 12 weeks old in Ringer solution that corresponded to 10% of the body weight (for example, 2.0 ml for a 20 g mouse, but not over 2.5 ml if the mouse was more than 25 g) within 6 s. Liver tumor experiments were conducted following institutional protocol in Massachusetts General Hospital. Mice were euthanized with CO2 followed by cervical dislocation. Tissues were harvested post euthanasia and analyze as indicated.

### Mouse brain homogenization and nuclear isolation

The mouse brain was harvested and rinsed with cold PBS. Homogenization buffer A (0.32 M sucrose, 1 mM NaHCO3, 0.25 mM CaCl2, 1 mM MgCl2, 50 mM Tris HCl, pH 7.5) supplied with halt proteinase and phosphatase inhibitors (78442, Invitrogen) was added as 2 ml buffer A per mouse brain. The tissue was homogenized with a Dounce homogenizer 20 times before being centrifuged at 1000g for 10 min. The supernatant was transferred to a new 15 ml falcon tube labeled as the cytoplasmic sample. For the pellet, 1 ml buffer A was used to resuspend it prior to further homogenization with a Dounce homogenizer 10 times. The sample was centrifuged (1000g, 10 minutes), and supernatant was collected and placed in a new tube to separate the pellet and supernatant.

The pellet was resuspended in 2 ml buffer B (0.32 M sucrose, 1 mM NaHCO3, 5 mM CaCl2, 3 mM MgCl2, 0.1 mM EDTA, 1 mM DTT, 0.1% NP40, 10 mM Tris HCl, pH 8.0), before being homogenized with a Dounce homogenizer 20 times. The sample was then centrifuged at 1000g for 10 minutes. The supernatant was discarded, and the pellet was resuspended with 5 ml buffer B and transferred into a 14 ml ultracentrifuge tube (331372, Beckman Coulter). 9 ml Buffer C (1.8 M sucrose, 3 mM MgCl2, 1 mM DTT, 10 mM Tris HCl, pH 8.0) was carefully added into the bottom of the tube. The samples were then ultracentrifuged at 24,000 rpm (106803.1x g RCF) for 1 hour prior to discarding of the supernatant (Thermo Scientific Sorvall WX floor ultra centrifuge and Surespin 630 swinging bucket rotor). Next, 1 ml cold PBS was added into the tube and kept on ice for 10 minutes. The pellet was then carefully resuspended and transferred into a new Eppendorf tube, prior to being centrifuged at 1000 g for 10 min. Finally, the supernatant was discarded; the pellet contains the nuclear fraction.

### Mouse brain immunoprecipitation

For the cytoplasm sample, 2X buffer D (300 mM NaCl, 2% NP40, 1% Sodium Deoxycholate, 2 mM EDTA, 2 mM EGTA, 100 mM Tris HCl, pH 7.5) was added and the sample was rotated for 1 hour. Next, the sample was centrifuged at 20,800g for 20 minutes and the supernatant was transferred into a new falcon tube. The sample was then centrifuged a second time under the same conditions, with the supernatant being collected. Of the sample (about 5 ml), 100 μl was saved as input and to test protein concentration. 25 μl GFP-trap beads (gmta-20 GFP-Trap ChromoTek GmbH) per mouse brain was added and the sample was incubated overnight. The next day, beads were washed three times (10 minutes each, 500 μl 1X buffer D). After the final wash, the protein was eluted at 70°C with 30 μl 1X LDS-PAGE sample buffer (NP0007). 3 μl of the IP sample was used for SDS-PAGE and Silver staining (LC6070, Invitrogen). The rest of the IP samples were subjected to mass-spectrometry analyses.

For the nuclear sample, 150 μl buffer E (Invitrogen (1861754) buffer with 1 mM MgCl2, 1 mM CaCl2, 10 mM ZnCl2, 1X halt proteinase and phosphatase inhibitor) was added. The sample was vortexed for 15 seconds at full speed, then incubated at 4°C for 30 minutes. Next, 1 μl benzonase (70746-4, Lot# 2705583, Millipore) was added and the sample was incubated for a further 30 minutes. Then 2X buffer D was added prior to an additional 30 minutes incubation on a rotator. The sample was then centrifuged at 20,800g for 10 minutes and the supernatant was transferred into a new Eppendorf tube. 10 μl GFP-trap beads were used per mouse brain to pull down the GFP-LC3 binding protein. The binding protein was eluted at 70°C with 20 μl 1X LDS-PAGE sample buffer. 2 μl of the IP sample was used for SDS-PAGE and Silver staining. The rest of the IP samples were subjected to mass-spectrometry analyses.

### Protein digestion and tandem-mass-tag (TMT) labeling of mouse brain

The analysis was performed with a previously optimized protocol^62–64^. For whole proteome profiling, quantified protein samples (1 mg in the lysis buffer with 8 M urea) for each TMT channel were proteolyzed with Lys-C (Wako, 1:100 w/w) at 21 °C for 2 h, diluted by 4-fold to reduce urea to 2 M for the addition of trypsin (Promega, 1:50 w/w) to continue the digestion at 21°C overnight. The insoluble debris was kept in the lysates for the recovery of insoluble proteins. The digestion was terminated by the addition of 1% trifluoroacetic acid. After centrifugation, the supernatant was desalted with the Sep-Pak C18 cartridge (Waters), and then dried by Speedvac. Each sample was resuspended in 50 mM HEPES (pH 8.5) for TMT labeling, and then mixed equally, followed by desalting for the subsequent fractionation. For whole proteome analysis alone, 0.1 mg protein per sample was used.

### Extensive two-dimensional liquid chromatography-tandem mass spectrometry (LC/LC-MS/MS) for mouse brain

The TMT labeled samples were fractionated by offline basic pH reverse phase LC, and each of these fractions was analyzed by the acidic pH reverse phase LC-MS/MS^65, 66^. The offline basic pH LC was performed with an XBridge C18 column (3.5 μm particle size, 4.6 mm × 25 cm, Waters), buffer A (10 mM ammonium formate, pH 8.0), buffer B (95% acetonitrile, 10 mM ammonium formate, pH 8.0), using a 2–3 h gradient of 15–35% buffer B^62^. In the acidic pH LC-MS/MS analysis, fractions were analyzed sequentially on a column (75 μm × 15–30 cm, 1.9 μm C18 resin from Dr. Maisch GmbH, 65 °C to reduce backpressure) coupled with a Fusion or Q Exactive HF Orbitrap mass spectrometer (Thermo Fisher Scientific). Peptides were analyzed with a 1–3 h gradient (buffer A: 0.2% formic acid, 5% DMSO; buffer B: buffer A plus 65% acetonitrile). For mass spectrometer settings, positive ion mode and data-dependent acquisition were applied with one full MS scan followed by a 20 MS/MS scans. MS1 scans were collected at a resolution of 60,000,1 × 10^6^ AGC and 50 ms maximal ion time; higher energy collision-induced dissociation (HCD) was set to 32–38% normalized collision energy; ∼ 1.0 m/z isolation window with 0.3 m/z offset was applied; MS2 spectra were acquired at a resolution of 60,000, fixed first mass of 120 m/z, 410–1600 m/z, 1 × 105 AGC, 100–150 ms maximal ion time, and ∼ 15 s of dynamic exclusion.

### Protein identification and quantification by the JUMP software suite for mouse brain

The bioinformatics processing of protein identification and quantification were carried out with the JUMP software suite^67–69^. In brief, MS/MS raw data were searched against a target-decoy database to estimate false discovery rate (FDR)^70^. We combined the downloaded Swiss-Prot, TrEMBL, and UCSC databases and removed redundancy (human: 83,955 entries) to create the database. Main search parameters were set at precursor and product ion mass tolerance (±15 ppm), full trypticity, maximal modification sites (n = 3), maximal missed cleavage (n = 2), static mass shift including carbamidomethyl modification (+ 57.02146 on Cys), TMT tags (+ 229.16293 on Lys and N-termini), and dynamic mass shift for oxidation (+ 15.99491 on Met). Peptide-spectrum matches (PSM) were filtered by mass accuracy, clustered by precursor ion charge, and the cutoffs of JUMP-based matching scores (J-score and ΔJn). The peptide was represented by the protein with the highest PSMs according to the rule of parsimony when one peptide was matched to multiple homologous proteins^71^. Protein quantification was performed based on the reporter ions from MS2 using our previously optimized method^65^.

### RNA-seq analysis

RNA quality was checked using Agilent Tapestation. RNA-seq libraries were prepared from total RNA using polyA selection followed by the NEBNext Ultra II Directional RNA library kit workflow (New England Biolabs). Sequencing was performed on the Illumina HiSeq 2500 instrument, resulting in approximately 30 million 50 bp reads per sample. Sequencing reads were mapped in a splice-aware fashion to the human reference transcriptome (hg19 assembly) using STAR^72^. Read counts over transcripts were calculated using HTSeq^73^ based on the Ensembl annotation for GRCh38/hg19 assembly. For the differential expression analysis, we used the EdgeR method^73, 74^ and defined differentially expressed genes (DEGs) based on the cutoffs of 2-fold change in expression value and false discovery rates (FDR) below 0.05. GO analysis was performed using EnrichR^36^. For heatmap visualization of SASP genes, the SASP factors curated previously were used as a reference^37^. The genes upregulated in HRasV12 samples (≥2 fold change compared to vector untreated samples) to a substantive expression level (RPKM ≥1 in HRasV12 samples) were used to generate heatmaps by an online tool^75^. Transcription factor predictions were performed using EnrichR^36^.

### ATAC-seq analysis

Sample preparations were done using established procedures^7647^. ATAC-seq libraries were sequenced on Illumina HiSeq 2500 instrument, resulting in approximately 40 million paired-end 50 bp reads per sample. Reads were mapped to the hg19 human reference genome using BWA^73, 74, 77^. The fragments with both ends unambiguously mapped to the genome that were longer than 100 bp were used in further analysis. Hotspot^78^ was used to detect significant peaks of read density with FDR cutoff of 0.05. Differentially accessible chromatin regions (DARs) were detected using the DiffBind package^79^ with FDR cutoff of 0.05. Pathway enrichment analysis was performed using EnrichR^36^. Heatmap visualization of ATAC-seq signal density at selected genes was based on RPKM values calculated over the regions including gene body and 1 kb flanks. In the comparison between vector HRasV12 and WSTF HRasV12 groups, cytokine and secreted factor genes with significant signal change (*P* < 0.05) were used. Genes with fold change >1.5 between HRasV12 and control were included in the heatmap.

### Fractionation and immunoprecipitation for primary human fibroblasts

Trypsinized cells were centrifuged at 500 g for 5 min at 4 °C. The cell pellets were lysed in buffer 1, pipetted up and down 10 times, and rotated for 30 min at 4 °C. Cells were then centrifuged at 500 g for 5 min at 4 °C; the collected supernatant is cytosolic fractionation. The pellets were further lysed with buffer 2, pipetted up and down 10 times, vortexed for 5 s, and rotated for 10 min at 4 °C, followed by centrifugation at 3000 g for 5 min at 4 °C. The collected supernatant is membrane fractionation. The pellets were lysed with buffer 3, pipetted up and down 10 times, vortexed for 10 s, and rotated for 1h at 4 °C. The product was centrifuged at 5000 g for 5 min at 4 °C. The collected supernatant is nuclear fractionation. Buffer 1: 150 mM NaCl, 50 mM HEPES, 25 ug/ml digitonin (Sigma 11024-24-1) with protease and phosphatase inhibitors (Roche); Buffer 2: 150 mM NaCl, 50 mM HEPES, 0.5% IGEPAL (Sigma I3021) with protease and phosphatase inhibitors (Roche); Buffer 3: 150 mM NaCl, 50 mM HEPES, 1% NP40 (Sigma 74285), benzonase (2 units/ml), 1 mM MgCl2, 0.1%SDS, 0.5% sodium deoxycholate with protease and phosphatase inhibitors (Roche). The nuclear fractionation was dialyzed using Amicon Ultra-0.5 Centrifugal Filter Unit (Millipore), followed by reconstitution and immunoprecipitation with buffer (20mM Tris HCl, pH 7.5, 137 mM NaCl, 1 mM CaCl2, 3 mM MgCl2, benzonase (2 units/ml), 0.5% NP40, with protease and phosphatase inhibitors (Roche)). Anti-HA magnetic beads (Thermo Fisher) were used for immunoprecipitation.

### Nuclear proteome analyses for primary human fibroblasts

For mass spectrometry of nuclear fractions and ATG8 immunoprecipitated, samples were alkylated and reduced using chloroacetamide, 20 mM at room temperature for 30 min and TCEP, 5 mM for 10 min at 55°C followed by TCA precipitation. TCA was added to eluates at final concentration of 20% and placed on ice at 4°C for at least an hour after which the samples were pelleted for 30 min. The pellets were washed three times using ice cold methanol. Dried pellets were resuspended in 50 μl, 200 mM EPPS, pH 8.0. Peptide digestion was carried out using LysC (Wako cat. # 129-02541, 0.25 μg) for 2 h at 37°C followed by trypsin (0.5 μg) overnight. Digested peptides were then labeled with 4 μl of TMT reagent (at 20 μg/μl stock) for 1 hr and the reaction was quenched using hydroxylamine at a final concentration of 0.5 % (wt/vol) for 20 min. The samples were then combined and dried in a vacuum centrifuge. This combined sample was then subjected to fractionation using the high pH reversed-phase peptide fractionation kit (Thermo Fisher) for a final of six fractions for the nuclear eluates. The dried fractions were processed by C18 stage tip desalting prior mass spectrometry.

Quantitative whole-cell proteomics of HRasV12 expressing cells was performed as per^80, 81^. After induction of senescence, cells were lysed in 8 M urea buffer (8 M urea, 1 M Tris pH 7.4, 5 M NaCl) containing protease and phosphatase inhibitors (Roche) followed by sonication at 4°C. The sonicated lysate was clarified by centrifugation at 13,000 rpm for 10 min at 4°C. Protein concentration was estimated by Bradford assay and 100 μg of total protein was used for each replicate. 100 μg of clarified lysate for each sample was then reduced and alkylated by the addition of 5 mM TCEP for 30 min at 37°C and 20 mM chloroacetamide for 20 min at room temperature, followed by methanol-chloroform precipitation at a 3:1 ratio to the lysate. The protein precipitate was washed thrice with ice cold methanol and resuspended in 200 mM HEPES pH 8.5 for protein digestion. Protein digestion was carried out by 1:100 protease to protein ratio of Lys-C for 2 h at 37°C followed by trypsin overnight. Each sample was then labeled with TMT reagent (TMT10 reagent, ThermoFisher Scientific, 90110) for 1 h, and the reaction was quenched with hydroxylamine at a final concentration of 0.3% w/v. 1% of each sample was mixed in a 1:1:1:1:1:1:1:1:1:1 ratio and a ‘ratio-check’ analysis using LC-MS/MS was performed to determine if the samples were present in equal ratios. Based on this result, the volumes of the remaining sample were adjusted and combined so as to maintain the 1:1:1:1:1:1:1:1:1:1 ratio. This combined sample was then dried to completeness using a vacuum centrifuge and acidified with 5% w/v formic acid. Digested peptides were cleaned up using C18 SPE (Sep-Pak, Waters) and separated using basic pH reversed-phase HPLC and pooled into 24 fractions. All 24 fractions were vacuum dried to completeness and subject to the C18 stage tip method prior to loading on the mass-spectrometer. Data was obtained using an Orbitrap fusion Lumos mass spectrometer linked with a Proxeon EASY-nLC 1200 LC pump. Peptides were separated on a 75 μM inner diameter microcapillary tube packed with 35 cm of Accucore C18 resin (2.6 μm, 100A, Thermo Fisher Scientific). The data were acquired using the MS3 method as outlined in previous study^82^.

Raw mass spectra obtained were processed using an in-house software pipeline as described previously^83, 84^. Values for protein quantification were exported and processed using Perseus to calculate Log fold changes and p values. The interaction maps were generated by Cytoscape software ^85^.

### Reagents and antibodies

All reagents were purchased from Sigma, unless otherwise stated. Odyssey ladder (Licor 928-60000). Primary antibodies used include: WSTF (Abcam ab51256), Lamin B1 (Abcam ab16048), phospho-Rb (Cell Signaling Technology 9308S), CyclinB1 (Cell Signaling Technology 12231S), phospho-S6 (Cell Signaling Technology 2215), GAPDH (Cell Signaling Technology 5174), IL8 (Abcam ab18672), p21 (Santa Cruz Biotechnology sc-271532), p16 (BD Biosciences G175-405), p-ATM (Abcam ab81292), IL6 (Cell Signaling Technology 12153), NF-κB p65 (Cell Signaling Technology 8242S and 6956S), phospho-p65 (Cell Signaling Technology 3033), IL1 alpha (Abcam ab18672), p38 MAPK (Cell Signaling Technology 9212), phospho-p38MAPK (Cell Signaling Technology 9211), phospho-ATF2 (Cell Signaling Technology 9221), phospho-p53 (Cell Signaling Technology 9284T), STING (Cell Signaling Technology 13647), Beta-tubulin (Sigma T8328-25UL), SNF2H (Abcam ab72499), HA-Tag (Sigma H3663-200UL), ACF1 (Abcam ab187670), Cyclin A2 (Cell Signaling Technology 4656), ATG7 (Cell Signaling Technology 8558S), BRG1 (Cell Signaling Technology 49360), ATG5 (Cell Signaling Technology 12994S), Histone 3 (H3) (Active Motif 39763), Histone H3K4me3 (Active Motif 39160), T7 Tag (MilliporeSigma 69522-3), H4K16ac (Millipore 07-329), Acetylated-Lysine (Cell Signaling Technology 9814S), IgG (Cell Signaling Technology 3900), Flag (Sigma F-1804), p65 K310Ac (Cell Signaling Technology 3045S), Ras (Cell Signaling Technology 14412), and CD45 (Biolegend 103101).

Secondary antibodies used for western blots include: Goat anti-Mouse IgG, IRDye® 680RD (Licor 926-68070), Goat anti-Mouse IgG, IRDye® 800RD (Licor 926-32210), Goat anti-Rabbit IgG, IRDye® 680CW (Licor 926-68071), Goat anti-Rabbit IgG, IRDye® 800CW (Licor 926-32211), Goat Anti-Mouse IgG (H + L)-HRP Conjugate (Bio Rad 1706516), Goat Anti-Rabbit IgG (H + L)-HRP Conjugate (Bio Rad 1706515), and Mouse Anti-Rabbit IgG (Light-Chain Specific) (Cell Signaling Technology 93702).

Secondary antibodies used for immunofluorescence (IF) include: Goat anti-Mouse IgG Alexa Fluor 488 (Thermo Fisher Scientific A-11001), Goat anti-Mouse IgG Alexa Fluor 532 (Thermo Fisher Scientific A-11002), Goat anti-Mouse IgG Alexa Fluor 555 (Thermo Fisher Scientific A-21422), Goat anti-Mouse IgG Alexa Fluor 647 (Thermo Fisher Scientific A-21236), Goat anti-Rabbit IgG, Alexa Fluor 488 (Thermo Fisher Scientific R-37116), Goat anti-Rabbit IgG Alexa Fluor 555 (Thermo Fisher Scientific A-21428), Goat anti-Rabbit IgG Alexa Fluor 647 (Thermo Fisher Scientific A-21446), Donkey anti-Rat IgG Alexa Fluor 647 (Thermo Fisher Scientific A-78947), Donkey anti-Goat IgG Alexa Fluor 555 (Thermo Fisher Scientific A-21432), and Donkey anti-Goat IgG Alexa Fluor 647 (Thermo Fisher Scientific A-21447).

### Retrovirus and lentivirus

Retroviral WZL-HRasV12 construct was transfected to the phoenix packaging cell line, and production of virus for stable expression was performed as previously described^21^. Lentiviral vectors were transfected with packaging plasmids to HEK293T cells, as described previously^21^. Viral supernatant was filtered through a 0.45-μm filter and mixed with trypsinized recipient cells. The infected cells were then selected with puromycin or hygromycin.

pLKO-shRNA constructs were purchased from Sigma. The following shRNA sequences were used: WSTF#1 (TRCN0000013341: GCAGATGACTTTGTTGGATAT) and WSTF#2 (TRCN0000013338: CCCACAACAAATCTAGCTCTA). Non-targeting control and ATG7 shRNA were described previously^6^. pLentiCRISPRv2 construct was used for CRISPR-mediated gene silencing. The following sequences were used: sgRNA targeting SNF2H #1 (CACCGAATCTTCAGTCAAATGACAA, AAACTTGTCATTTGACTGAAGATTC). sgRNA targeting SNF2H #2 (CACCGATAGCCTGAAGATCTACTTG, AAACCAAGTAGATCTTCAGGCTATC). Control sgRNA was described previously^6, 7^.

### Plasmids

GST, GST–LC3A, B, C, GABARAP, GABARAPL1 and GABARAPL2 were described elsewhere^86^. HA-tagged LC3A, B, C, GABARAP, GABARAPL1 and GABARAPL2 cDNA were described previously^15^ and were cloned into lentiviral vectors for stable expression. GABARAP truncations/mutations were made from GST-GABARAP. WSTF and SNF2H cDNA were cloned into the *in vitro* translation (IVT) vectors for *in vitro* translation (1-Step IVT Systems, Thermo Fisher) and lentiviral/retroviral vectors for stable expression. WSTF truncations/mutations were made from plenti-HA-WSTF. All new constructs in this study were verified by DNA sequencing. T7-tagged p65 WT and mutants were purchased from Addgene: T7-RelA (#21984), T7-RelA(3KR) (#23251), T7-RelA(K310R) (#23250), T7-RelA(K221R) (#23248), T7-RelA(K218R) (#23247). HA-p300 was purchased from Addgene (#89094).

### Immunoblotting

Western blotting was described previously^21^ with slight modifications. Cells were lysed in buffer containing 20 mM Tris, pH 7.5, 137 mM NaCl, 1 mM MgCl2, 1 mM CaCl2, 1% NP-40, supplemented with 1:100 Halt protease and phosphatase inhibitor cocktail (Thermo) and benzonase (Novagen) at 12.5 U/mL. The lysates were rotated at 4 °C for 30 min and boiled at 95 °C in the presence of 1% SDS. The resulting supernatants were subjected to electrophoresis using NuPAGE Bis-Tris precast gels (Thermo). After transferring to nitrocellulose membrane, 5% milk in TBS was used to block the membrane at room temperature for 1 h. Primary antibodies were diluted in 5% BSA in TBS supplemented with 0.1% Tween 20 (TBST) and incubated at 4 °C overnight. The membrane was washed 3 times with TBST, each for 10 min, followed by incubation of secondary antibodies at room temperature for 1 h, in 5% milk in TBST. The membrane was washed again 3 times and imaged by film or by an Odyssey imager (Licor Odyssey CLx 2000).

### Immunofluorescence

Immunofluorescence was performed as described previously^21^. Briefly, cells were fixed in 4% paraformaldehyde in PBS for 30 min at room temperature. Cells were washed twice with PBS, and permeabilized with 0.5% Triton X-100 in PBS for 10 min. After washing twice, cells were blocked with 10% BSA in PBS for 1 h at room temperature. Cells were then incubated with primary antibodies in 5% BSA in PBS supplemented with 0.1% Tween 20 (PBST) overnight at 4 °C. The next day, the cells were washed four times with PBST, each for 10 min, followed by incubation with Alexa Fluor-conjugated secondary antibody (Thermo), in 5% BSA/PBST for 1 h at room temperature. The cells were then washed four times with PBST, incubated with 1 μg/mL DAPI in PBS for 10 min, and washed twice with PBS. The slides were mounted with ProLong Diamond (Thermo) and imaged with a Leica TCS SP8 fluorescent confocal microscope. Quantification of % positive cells was done under identical microscopy settings between samples. Over 200 cells from 4 randomly selected fields were analyzed.

### *In vitro* translation

*In vitro* translation was performed using the One Step *In Vitro* Translation Kit (Thermo Scientific). Target proteins were cloned into pT7CFE1-NHA vector (with N-terminal HA tag) and translated *in vitro* at 30 °C for 6 hours.

The other *in vitro* translation was done in the presence of radioactive [^35^S]methionine using the TNT T7 Reticulocyte Lysate System (Promega 14610). *In vitro*–translated proteins were then precleared by incubation with empty glutathione Sepharose beads in NETN buffer (50 mM Tris, pH 8.0, 150 mM NaCl, 1 mM EDTA, and 0.5% NP-40) supplemented with protease inhibitors for 30 min at 4°C to remove nonspecific binding. The precleared lysates were incubated with the GST fusion protein–loaded beads for subsequent GST pulldown experiments.

### Bacterial expression and GST pulldown

cDNA of target proteins were cloned into GST fusion expression vectors. GST-tagged vectors were transformed in BL21-CodonPlus *Escherichia coli,* then expressed and purified with glutathione beads (Thermo Scientific #78602) at 4 °C for 2 h and washed four times with the wash buffer containing 50 mM Tris, pH 7.5, 150 mM NaCl, 1% Triton X-100, 1 mM DTT, supplemented with 100 μM PMSF. The purified proteins or *in vitro* translated proteins were diluted in binding buffer (20 mM Tris, pH 7.5, 137 mM NaCl, 1 mM MgCl2, 1 mM CaCl2, 1% NP-40, supplemented with 1:1000 Halt Protease inhibitor cocktail) and then pre-cleared with GST at 4 °C for 1 h. The resulting supernatant was then subjected to GST pull-down with GST or GST fusion proteins. The product was washed four times with the wash buffer and boiled with NuPAGE loading dye for immunoblotting analysis. WSTF peptides were purchased from Genscript and purified by Pierce ™ C-18 Spin Columns (Thermo Fisher 89870).

### Immunohistochemistry

Mouse liver was buried in O.C.T. Compound (Thermo Fisher 23-730-571) and frozen on dry ice. The frozen tissues were sectioned into desired thickness using a cryostat (35 µm) (Leica). The tissue sections were placed onto glass slides suitable for immunohistochemistry, and oaked in 4% PFA for 20 min. The slides were rinsed twice gently with PBS, followed by permeabilization and blocking using filtered (by a 0.45 uM syringe filter) 10% BSA and 0.5% Triton X100 in PBS for 2 h. A circle was drawn surrounding the tissue sample using a hydrophobic pen. Primary antibody was added (1:100-500) into the circle with filtered 10% BSA in PBST (Tween-20 0.1%) and incubated in a humid box overnight at 4 degrees. The slides were gently rinsed 3 times with PBST on a shaker for 10 min each. 2nd fluorescent antibodies (1:500) were incubated for 2 h at room temperature on a shaker with gentle shaking. The slides were then gently rinsed 4 times with PBST for 10 min each, followed by incubation with DAPI (1 ug/ml in PBS) for 20 min. The samples were then gently rinsed twice with PBS for 10 min each, and mounted with sufficient ProLong™ Diamond Antifade Mountant (Thermo Fisher P36961). The slides were dried in a dark place overnight and imaged by a confocal microscopy.

### RT-qPCR

mRNA was extracted from cells or tissues using RNeasy Mini Kit (Qiagen), with a DNase I (Qiagen) digestion step to minimize genomic DNA contamination. Reverse transcription (RT) was done using High-Capacity RNA-to-cDNA Kit (Thermo), and then quantitative PCR (qPCR) was performed using a qPCR machine (BioRad CFX 384 Real-Time System). Results were normalized to Lamin A/C for both human cells and mouse cells. The following primers were used for RT-qPCR of human cells: WSTF: GCCAAAGGCACGCAGAAGAT, GGCTGCAGGAGGCAGATGTT; LaminB1: CTCGTCGCATGCTGACAGAC, GATCCCTTATTTCCGCCATCT; p16: CCAACGCACCGAATAGTTACG, CCATCATCATGACCTGGATCG; IL6: CACCGGGAACGAAAGAGAAG, TCATAGCTGGGCTCCTGGAG; p21: CTCAGGGGAGCAGGCTGAAG, AGAAGATCAGCCGGCGTTTG; IL8: ACATGACTTCCAAGCTGGCC, CAAATCAGGAAGGCTGCC IL1α: AGTGCTGCTGAAGGAGATGCCTGA, CCCCTGCCAAGCACACCCAGTA IL1β: CTCTCTCCTTTCAGGGCCAA, GAGAGGCCTGGCTCAACAAA. Lamin A/C: AGCTGAAAGCGCGCAATACC, GGCCTCCTTGGAGTTCAGCA; SNF2H: TGATGCGTCACCTGGAAAGC, GCCCGGTCAGTTTGCATTTT.

For mouse tissues, RNeasy Mini Kit (Qiagen) was used to extract the RNA. For mouse liver NRasV12 related RT–qPCR, three pieces of liver from the same mouse were combined as one sample (n = 1), and the mRNA extraction and reverse transcription were performed as previously described21. The results of SASP/immune cell factors were normalized to the value of NRasV12 as an internal control for NRas abundance. The following primers were used for RT-qPCR of mouse cells/tissues: WSTF: TGCGGGAAAAAGCCAAAGAA, CAGCCTGAATGCTGGGAGGT; IL-1α: TTCAAGGAGAGCCGGGTGAC, TGCTGATCTGGGTTGGATGG; IL1β: TCGCAGCAGCACATCAACAA, GCTGCCACAGCTTCTCCACA; IL6: CCGTGTGGTTACATCTACCCT, CGTGGTTCTGTTGATGACAGTG; CXCL1: CCATGGCTGGGATTCACCTC, CCAAGGGAGCTTCAGGGTCA; NRasV12: CCTCAGCCAAGACCAGACAGG, CATCACCACACATGGCAATCC; GAPDH: TGCATCCTGCACCACCAACT, ACGCCACAGCTTTCCAGAGG; CD45: CTGGAAGGCCTGGAAGCAGA, TGTGCCTCCACTTGCACCAT; CD3: CGAGGAACCGGTGCTGGTAG, CTGGGTTGGGAACAGGTGGT; Klrd1: GCCTTCTTCAGCCCCAATCC, TGTGCCATCCTCCCATAGCC. Lamin A/C: TTCCCTGGAGACCGAGAACG, CACCTCTCGGCTGACCACCT.

### Conditioned media and cytokine arrays

Control or senescent cells were cultured in regular media for 2 to 3 days before use. The media were then collected and filtered with a 0.45 µm PVDF filter (Millipore) to remove cells and debris. The resulting supernatant was used for immunoblotting. The amounts of media used for immunoblotting were quantified based on the protein concentrations of cell lysates, and media corresponding to equal amounts of total cellular proteins were loaded into protein gels. Cytokine arrays were performed as described previously^21^. The serum-free conditioned media were analyzed by a cytokine array (RayBiotech, Human Cytokine Array C3) following the manufacturer’s guidelines. The intensities of array dots were quantified with Fiji and normalized against the positive controls on the blots.

### Data mining from published resources

Quantitative proteomic data of the Cancer Cell Line Encyclopedia (CCLE) were based on a previous report^49^. Cancer types from solid tissues were used. For box plots displayed in this study, the central rectangle spans a range from the first quartile to the third quartile, also called the Interquartile Range. A line inside the rectangle shows the median. The whiskers are drawn down to the 10th percentile and up to the 90th. Points below and above the whiskers are drawn as individual dots. Outliers were defined as data points that were either 1.5 × interquartile range or more above the third quartile, or 1.5 × interquartile range or more below the first quartile. Unpaired t test with Welch’s correction (do not assume equal SDs) was used to compute statistical significance. P values less than 0.05 were considered significant.

### Statistical analyses

Unpaired two-tailed Student’s t-test was used for comparison between two groups. One-way ANOVA coupled with Tukey’s post hoc test was used for comparisons over two groups. Two-way ANOVA coupled with post hoc test was used for comparisons whereas there are two independent variables. Significance was considered when p value was less than 0.05.

### Data availability

RNA-seq and ATAC-seq data have been deposited in the NCBI Gene Expression Omnibus (GEO) database under accession number GSE214410. Other original data are available upon reasonable request.

## Extended Data Figure Legends

**Extended data figure 1.**
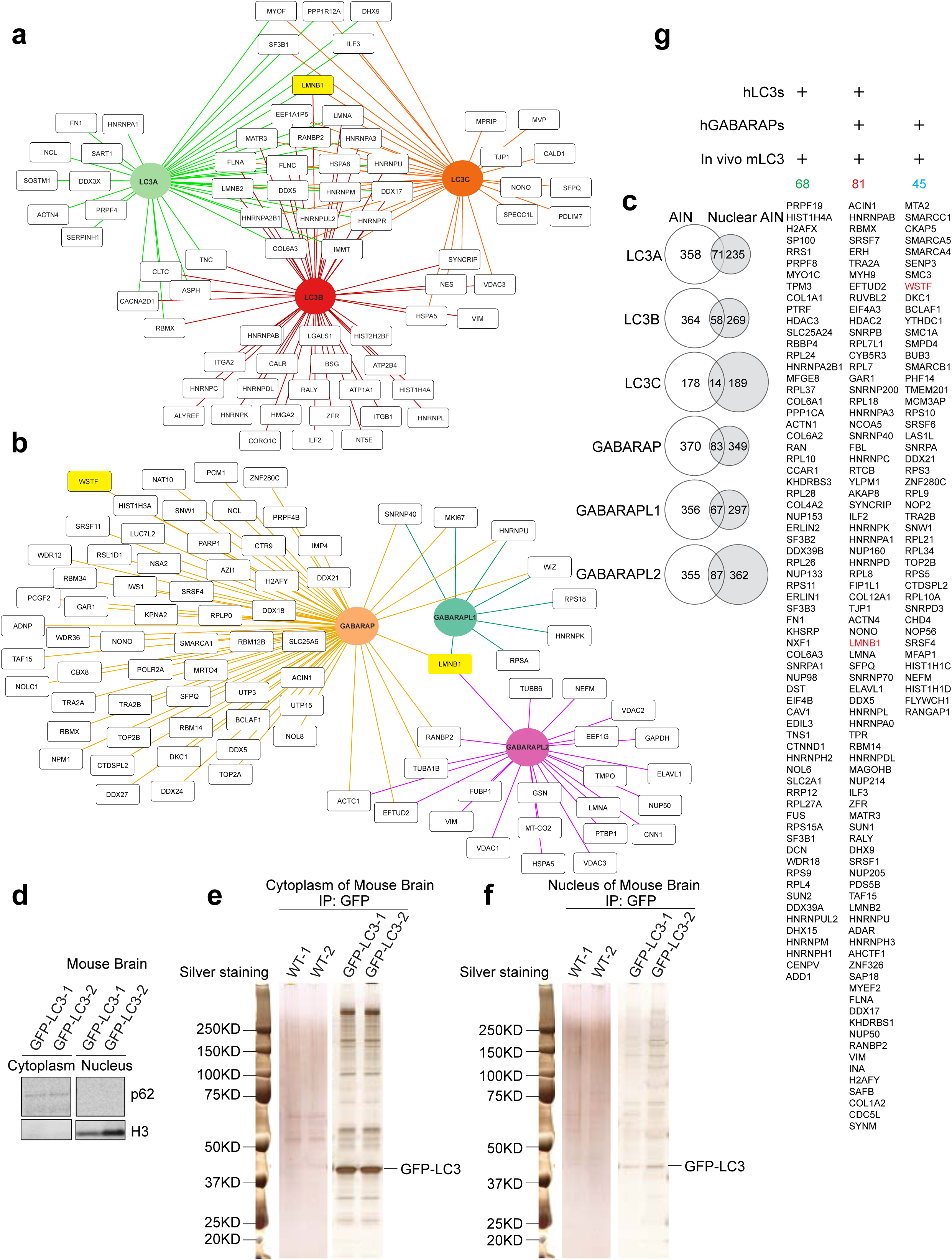
Nuclear ATG8 interactome. **a**, Interaction maps showing the nuclear binding partners of human MAP1LC3 isoforms (MAP1LC3A, MAP1LC3B and MAP1LC3C) from the proteomic analysis, generated by Cytoscape software. **b,** Interaction maps showing the nuclear binding partners of human GABARAP isoforms (GABARAP, GABARAPL1 and GABARAPL2) from proteomic analysis. **c,** Venn diagrams showing the overlap of autophagy interaction network (AIN) from a previous study versus the ATG8 nuclear interaction network from this study. **d,** Western blotting showing the fractionation of cytoplasm and nucleus from the brain of GFP-LC3B transgenic mice. **e and f,** The cytoplasm (**e**) and nucleus (**f**) fractions of the mouse brain were subjected to GFP IP. Silver-stained protein gels are shown. **g,** A list showing the overlap of ATG8 interacted nuclear proteins among the human LC3s (MAP1LC3A, MAP1LC3B and MAP1LC3C), human GABARAPs (GABARAP, GABARAPL1 and GABARAPL2), and *in vivo* mouse LC3.

**Extended data figure 2.**
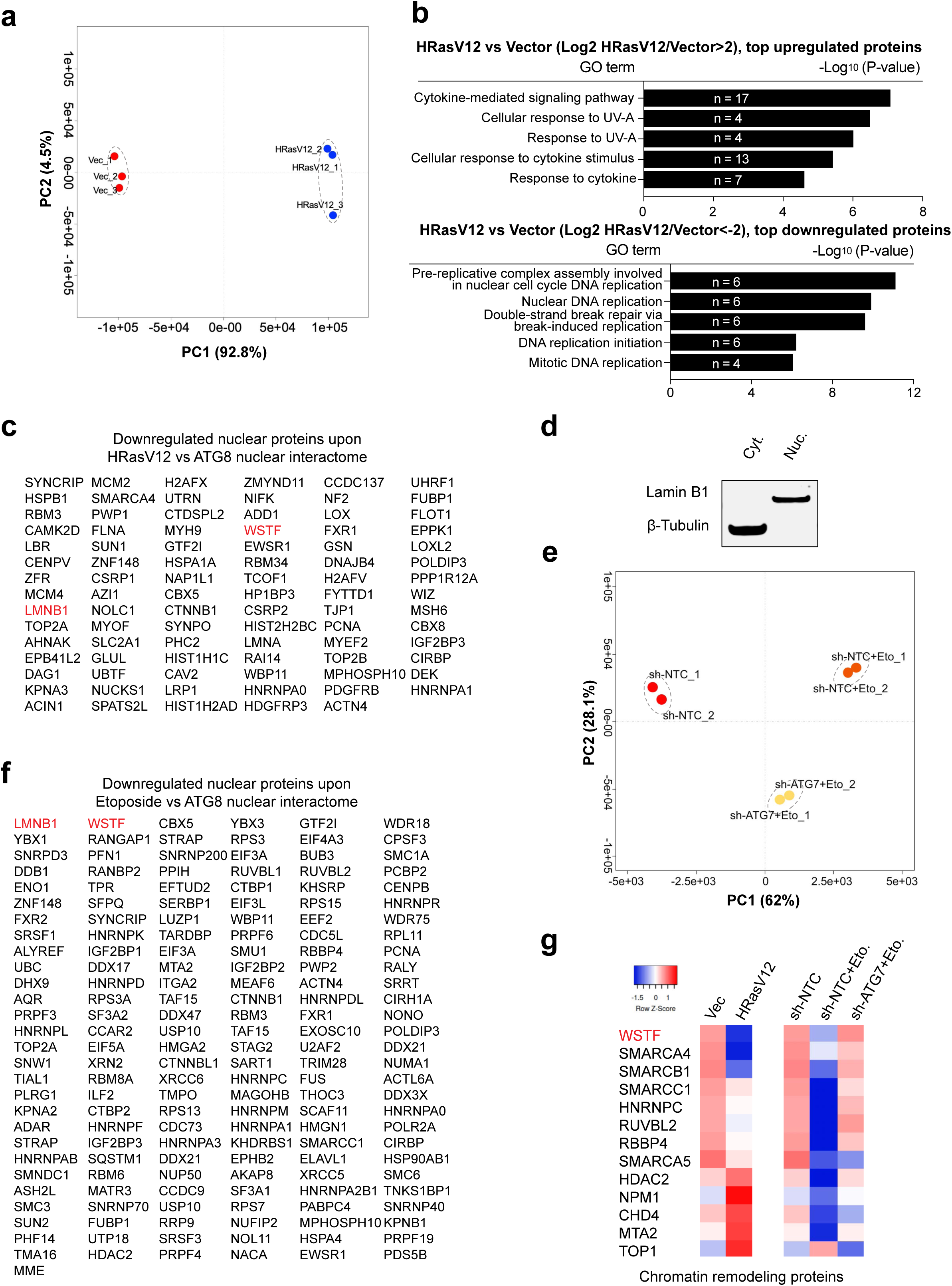
Proteomic analyses of nuclear proteomes in oncogene-induced senescence and etoposide-induced senescence. **a,** PCA analysis of the whole-cell proteomes from control and oncogene-induced senescent cells. **b,** GO analysis showing the top biological processes for upregulated and downregulated proteins in HRasV12-induced senescent cells. **c**, Related to Fig. 2e, a list showing the overlapping proteins between downregulated nuclear proteins upon HRasV12 and ATG8 nuclear interactome in human fibroblasts. **d**, Related to Fig. 2f, western blotting showing the fractionation of cytoplasm and nucleus. **e,** PCA analysis of the nuclear proteomes from untreated, etoposide-treated, and etoposide-treated IMR90 cells with ATG7 knockdown. **f,** Related to Fig. 2j, a list showing the overlapping proteins between downregulated nuclear proteins upon etoposide and ATG8 nuclear interactome in human fibroblasts. **g,** Heatmap presentation of the protein levels of chromatin remodeling proteins upon HRasV12 and etoposide-induced senescence with sh-NTC or sh-ATG7.

**Extended data figure 3.**
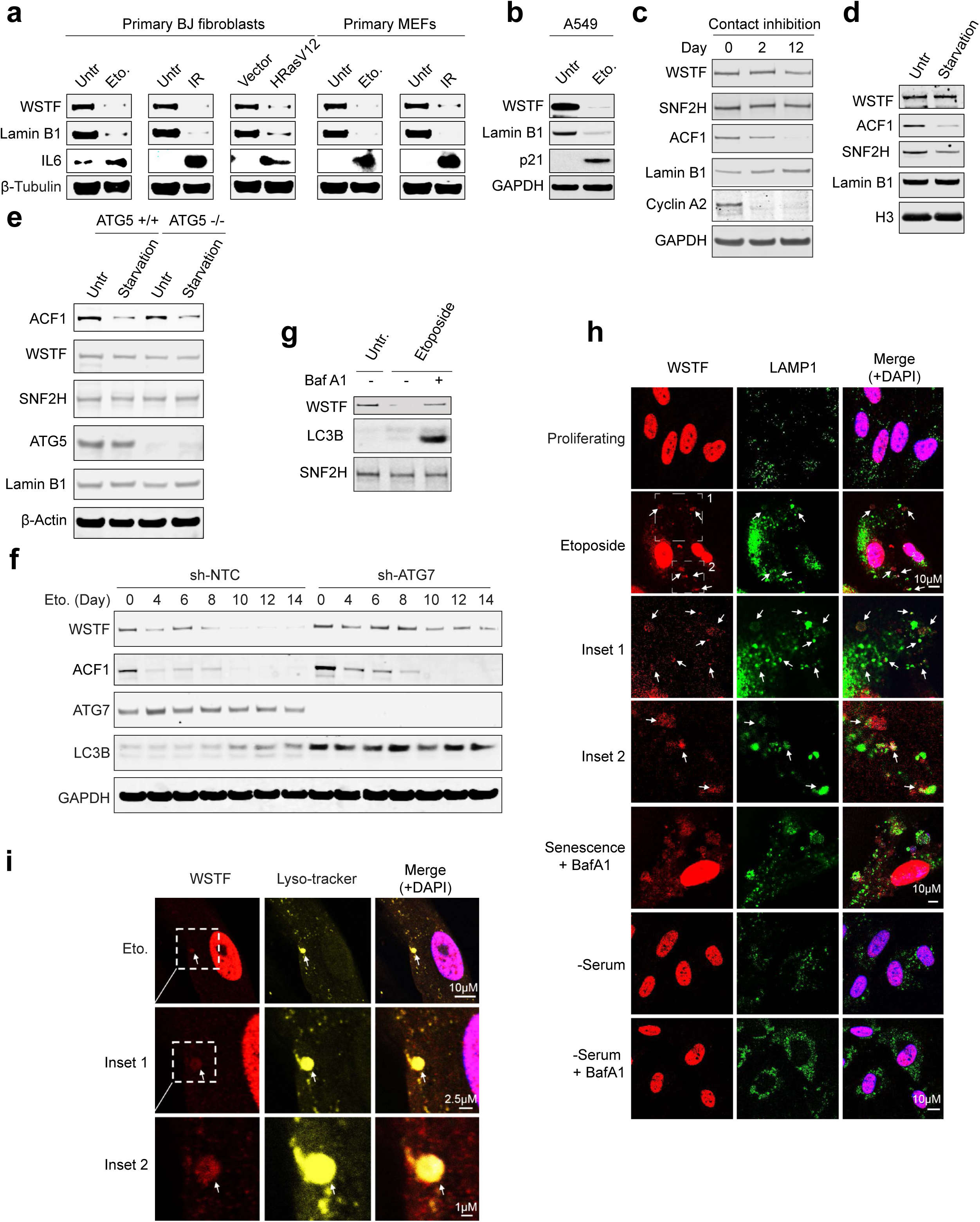
WSTF is degraded by autophagy during cellular senescence. **a**, Primary BJ fibroblasts and primary MEFs were induced to senescence using various means, and were analyzed by western blotting using indicated antibodies. **b**, A549 cells were treated with etoposide to induce senescence, and then analyzed by western blotting using indicated antibodies. **c**, IMR90 cells were induced to quiescence by contact inhibition, and were harvested at indicated days and analyzed by western blotting. **d**, IMR90 cells were treated by amino acid starvation for 24 hours and analyzed by western blotting. **e**, Wild-type and ATG5 knockout MEFs were treated by amino acid starvation for 24 hours and analyzed by western blotting. **f**, IMR90 cells stably expressing sh-NTC or sh-ATG7 were treated by etoposide and harvested at indicated days, followed by western blotting analyses. **g**, IMR90 cells were induced to senescence by etoposide, and bafilomycin A was added to the media of senescent cells, followed by western blotting analyses. **h**, IMR90 cells were left untreated or treated with etoposide to induce senescence, or serum starvation for 24 hours, with or without bafilomycin A1 added for 24 hours as indicated, and were stained with WSTF and LAMP1 antibodies, followed by imaging under a confocal microscopy. Representative images and scale bars are shown. **i**, IMR90 cells were induced to senescence by etoposide, then stained with Lyso-tracker and WSTF antibody, followed by imaging under a confocal microscopy. Representative images and scale bars are shown.

**Extended data figure 4.**
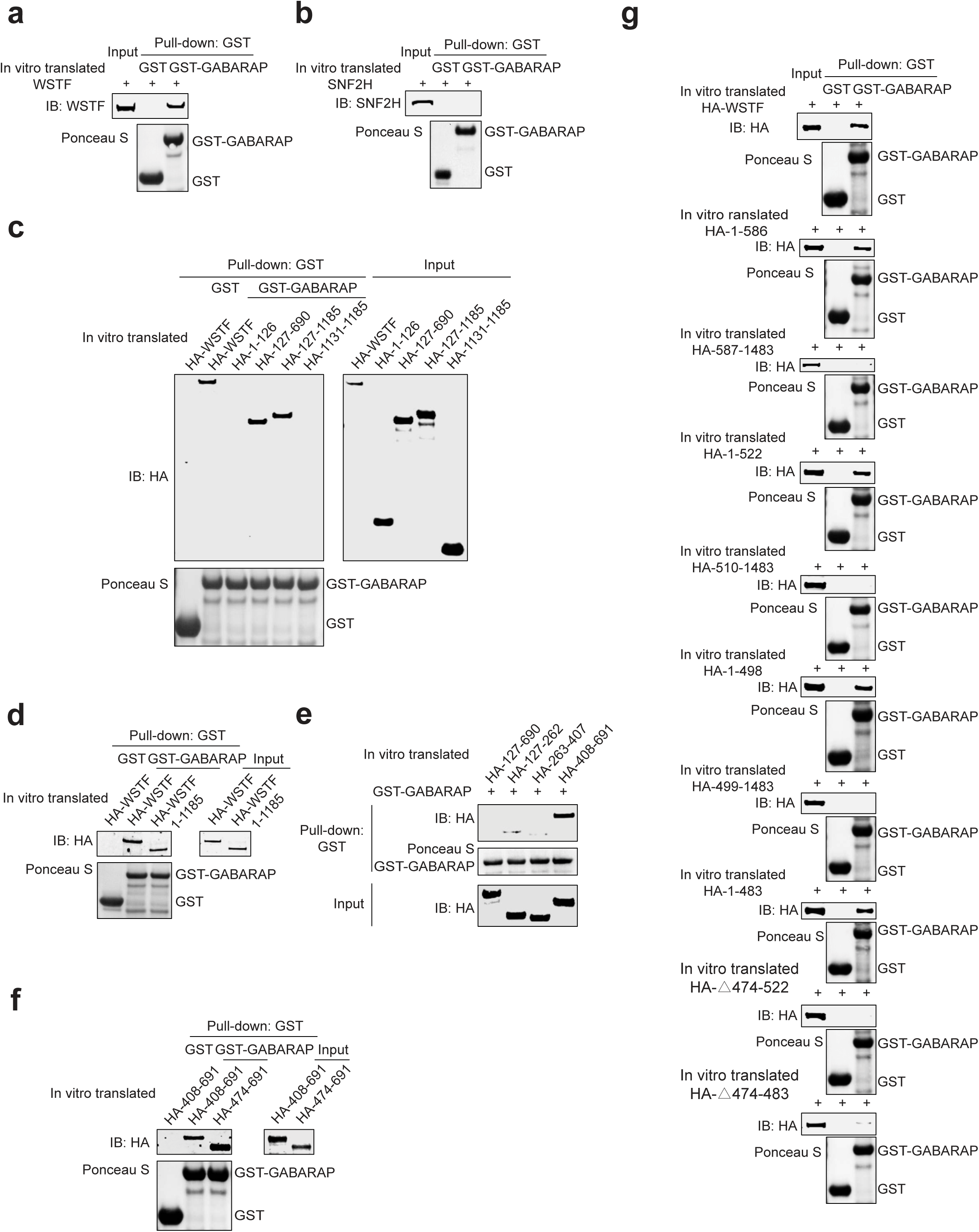
Characterization of GABARAP and WSTF association. **a**, *In vitro* translated WSTF were subjected to GST-tagged GABARAP pulldown and analyzed by western blotting. **b**, *In vitro* translated SNF2H were subjected to GST-tagged GABARAP pulldown and analyzed by western blotting. **c to g**, *In vitro* translated WSTF wild-type and mutants as shown were subjected to GST-tagged GABARAP pulldown followed by western blotting analyses.

**Extended data figure 5.**
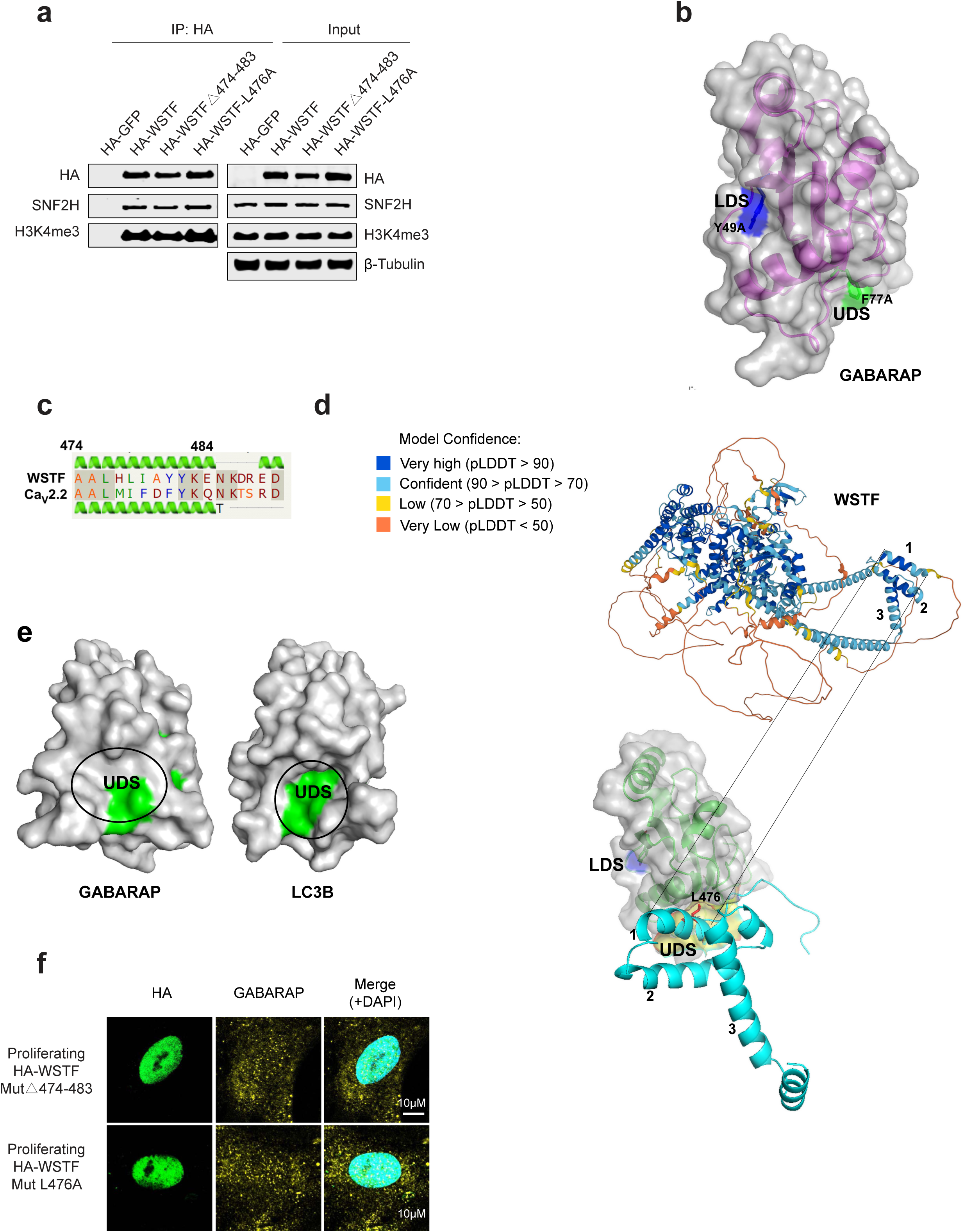
Molecular mechanisms for GABARAP and WSTF association. **a**, WSTF wild-type and mutants as shown were subjected to HA IP and analyzed by western blotting. **b**, Surface model (grey) and secondary structure of GABARAP (PDB code:1GNU) showing the locations of the LDS and UDS and the mutants compromising the respective sites indicated. **c**, PHYRE2 prediction of the hydrophobic helix of WSTF aligned with the published structure of Cav2.2. **d**, Upper part, Alphafold2 prediction of WSTF protein structure (UniProt:Q9UIG0). The helix 1 predicted by PHYRE2 is here predicted with high confidence. Lower part, model with helix1 and the exposed L476 docked into the UDS pocket of GABARAP. The region of the Alphafold2 model encompassing amino acids 460 to 550 of WSTF with the helices 1,2 and 3 was used as input with the solved structure of GABARAP (PDB code: 1GNU) for automatic protein-protein docking on the ClusPro web server. WSTF region 460-550 is shown in magenta, the L476 residue in red, and the contact surface with the UDS of GABARAP shown in yellow. Helices 1-3 are indicated. **e**, Surface models of structures of GABARAP (PDB code: 1GNU) and LC3B (PDB code: 5D94) showing the UDS pockets with some directly aligned residues (ALFFFV for GABARAP and AFFLLV for LC3B) in green. **f**, Related to Fig. 4 **p,q**, Proliferating IMR90 cells expressing WSTF mutants were stained with HA and GABARAP antibodies, followed by imaging under a confocal microscopy. Representative images and scale bars are shown.

**Extended data figure 6.**
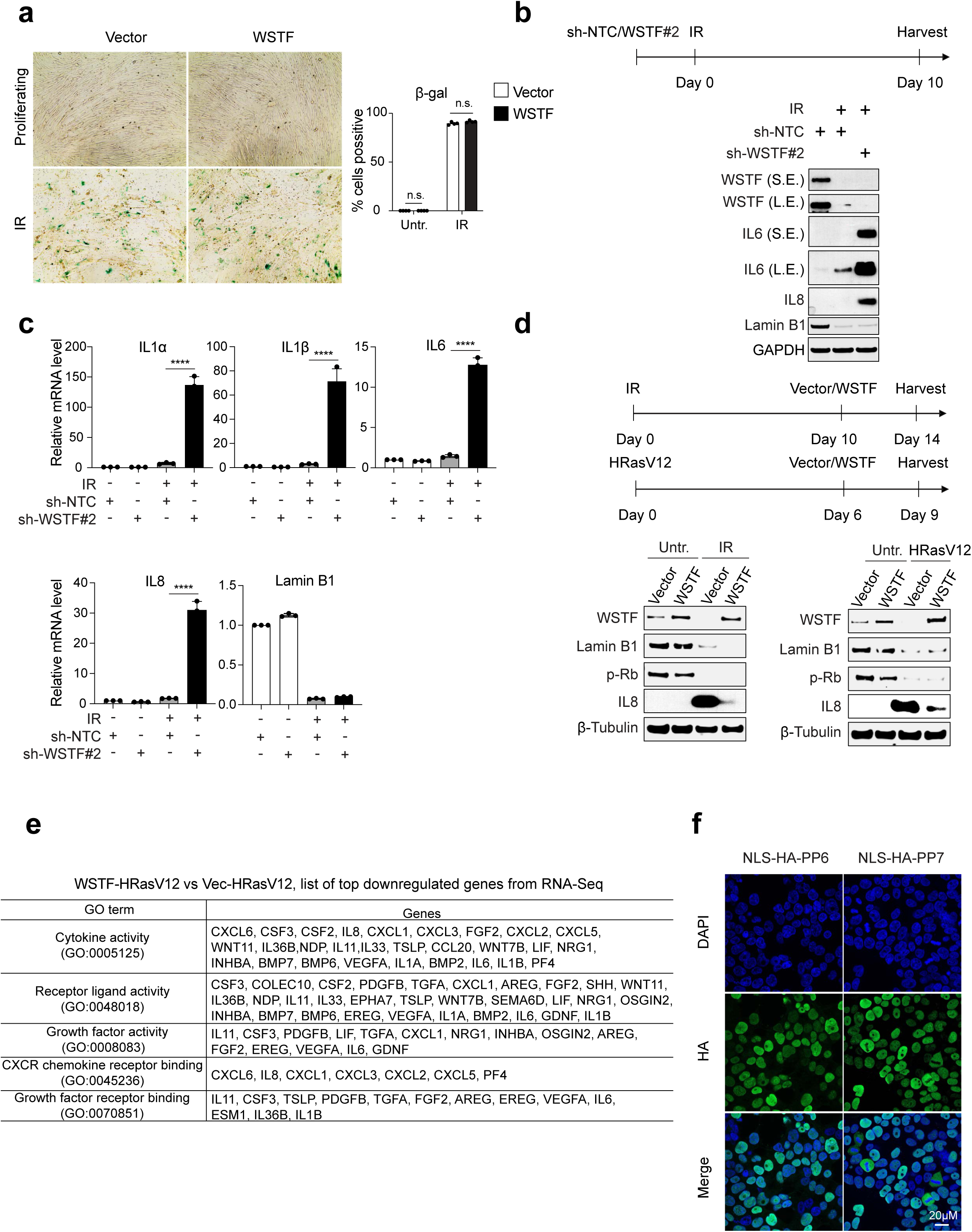
WSTF is a negative regulator of the SASP program. **a**, β-gal staining of IMR90 cells with vector control or WSTF overexpression. The cells were left untreated or subjected to IR-induced senescence and fixed at day 14. Bar graphs showing the quantification of the percentage of β-gal positive cells from 4 randomly selected fields. n.s.: non-significant. **b and c**, IMR90 cells were stably expressed with sh-NTC or sh-WSTF #2 hairpin, and treated with or without IR as indicated in the scheme, followed by analyses of western blotting (**b**) and RT-qPCR (**c**). **c**, RT-qPCR analyses of ILα, IL1β, IL6, IL8, and Lamin B1. Results were normalized to Lamin A/C and presented as mean values with s.d.; n=3; **** P < 0.0001; unpaired two-tailed Student’s t-test. **d**, IMR90 cells were treated with IR or HRasV12 to establish senescence, and then were infected with lentivirus encoding vector or WSTF, followed by western blotting analyses. **e**, Related to Fig. 5j, a list showing the genes downregulated by WSTF upon HRasV12-induced senescence, from RNA-seq studies. **f**, HEK293T cells were transfected with NLS-HA-PP6 and NLS-HA-PP7 constructs, and then stained with DAPI and HA antibody, followed by imaging under a confocal microscopy. Representative images are shown.

**Extended data figure 7.**
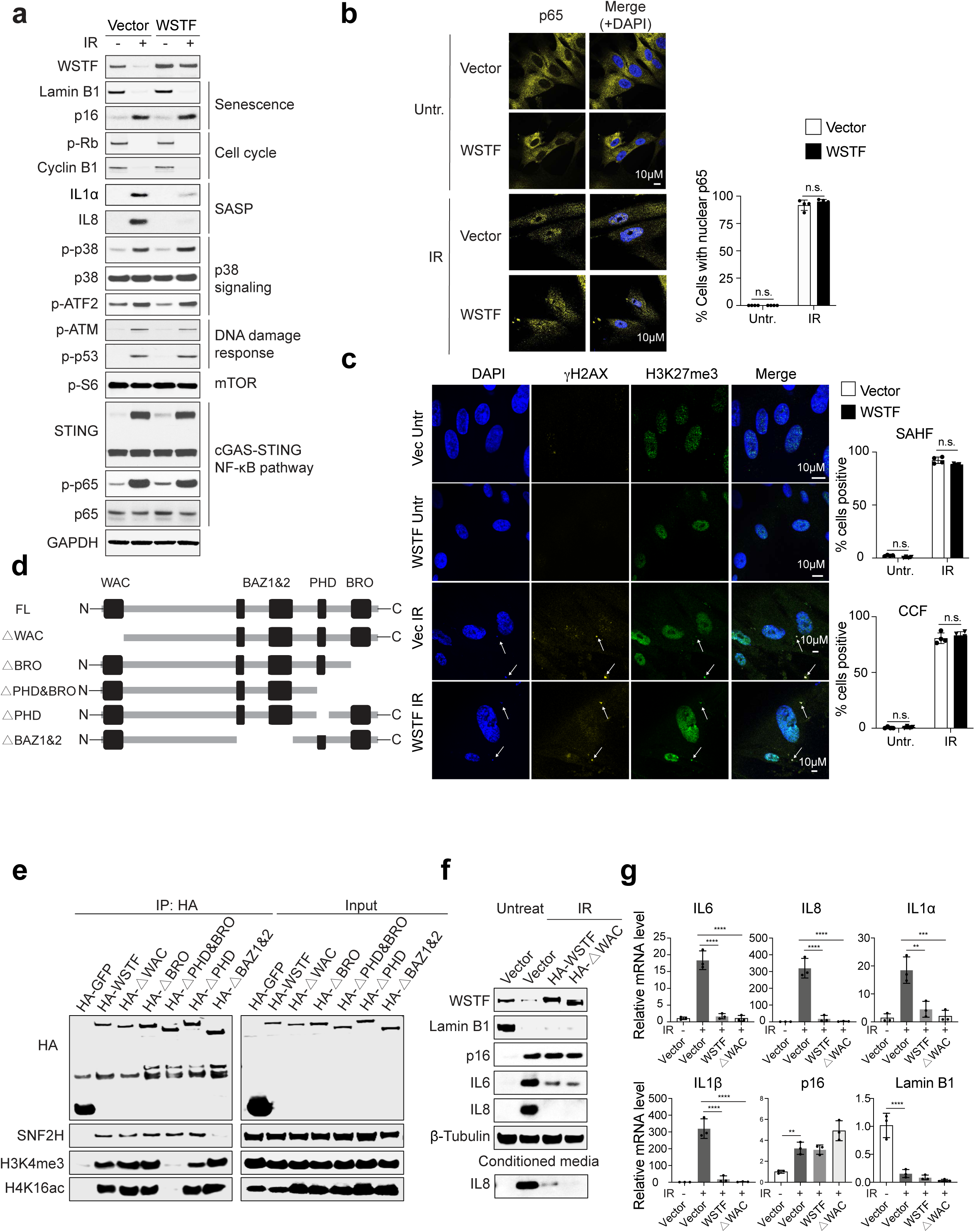
Molecular mechanisms underlying WSTF regulation of the SASP program. **a**, IMR90 cells stably expressing vector control or WSTF were left untreated or treated with IR (20 Gy), and were harvested at day 10. The lysates were analyzed by immunoblotting using indicated antibodies. **b**, Vector and WSTF stably expressing IMR90 cells were left untreated or induced to senescence with IR, then stained with a p65 antibody, followed by imaging under a confocal microscopy. Representative images are shown. Bar graphs showing the quantification of the percentage of nuclear p65 positive cells in vector and WSTF cells with or without IR treatment. Results shown are the mean values from four randomly selected fields with over 200 cells. Error bars: s.d.; n.s.: non-significant. **c,** IMR90 cells were left untreated or induced to senescence with IR, and stained with γH2AX and H3K27me3 antibodies, followed by imaging under a confocal microscopy. Representative images are shown. Arrows indicate CCF. Bar graphs showing the quantification of the percentage of SAHF and CCF positive cells in vector and WSTF cells with or without IR treatment. Results shown are the mean values from four randomly selected fields with over 200 cells. Error bars: s.d.; n.s.: non-significant. **d**, Related to Fig. 6a, schematic illustrations of WSTF full-length and truncations. **e**, HEK293T cells were transfected with HA-tagged wild-type or mutant WSTF constructs, and then subjected to HA immunoprecipitation and immunoblotting with indicated antibodies. **f and g**, IMR90 cells stably expressing WSTF wild-type or mutant were treated with or without IR, then the cell lysates and conditioned media were analyzed by western blotting (**f**) or RT-qPCR (**g**). **g**, RT-qPCR analyses of IL6, IL8, ILα, IL1β, p16 and Lamin B1. Results were normalized to Lamin A/C and presented as mean values with s.d.; n=3; ** P < 0.01; *** P < 0.001; **** P < 0.0001; one-way ANOVA coupled with Tukey’s post hoc test.

**Extended data figure 8.**
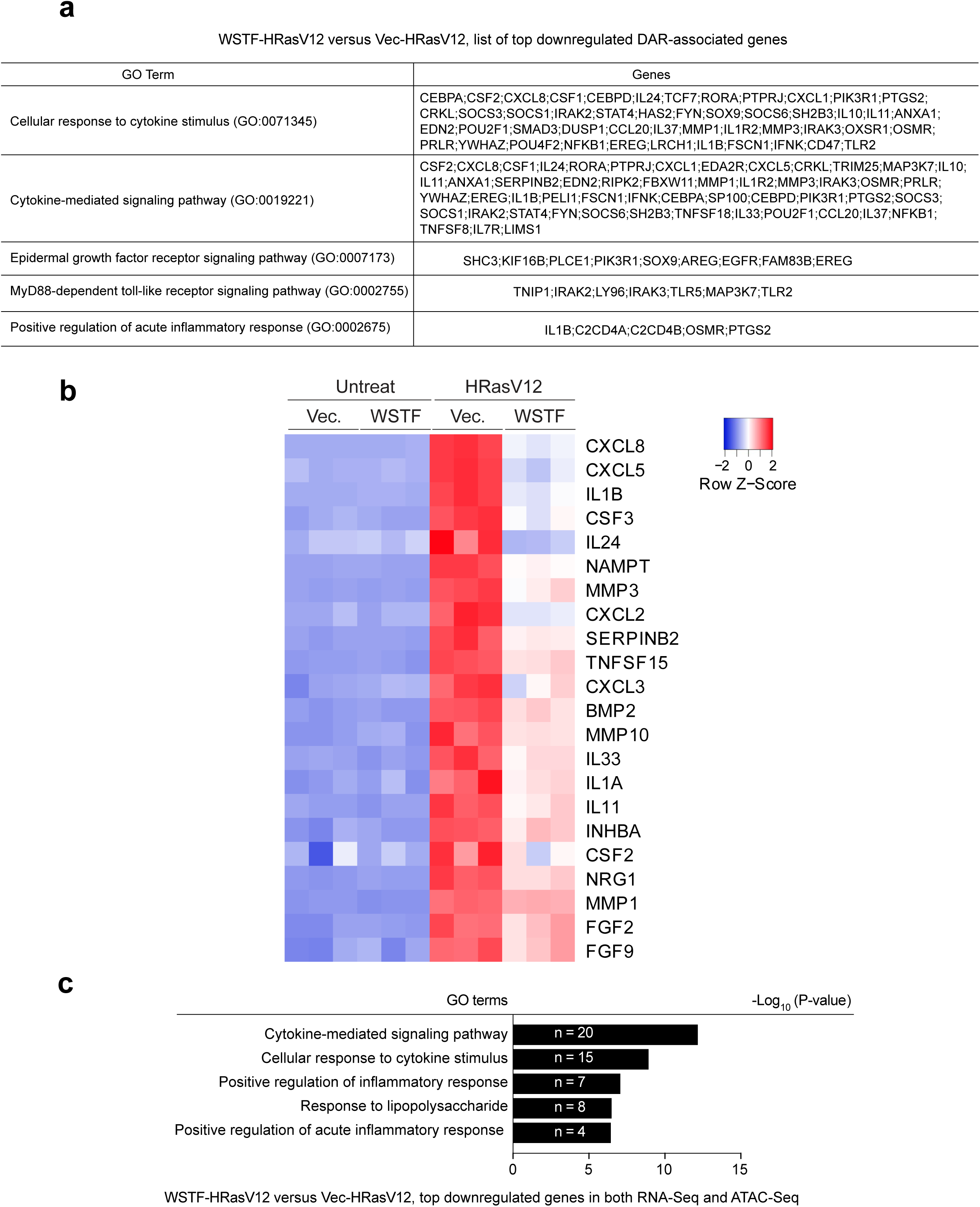
WSTF reduces chromatin accessibility at SASP genes. **a**, Related to Fig. 6j, list of genes corresponding to DARs whose chromatin accessibility was decreased by WSTF in HRasV12-induced senescence. **b**, Heatmap of the ATAC-seq signal density (RPKM) at TSS-proximal region, gene body, and TES-proximal region of inflammatory genes from. **c**, DEGs downregulated by WSTF according to RNA-seq were overlapped with genes of DARs with chromatin accessibility decreased by WSTF based on ATAC-seq. The GO terms enriched among these genes are shown with the corresponding numbers of genes and P values.

**Extended data figure 9.**
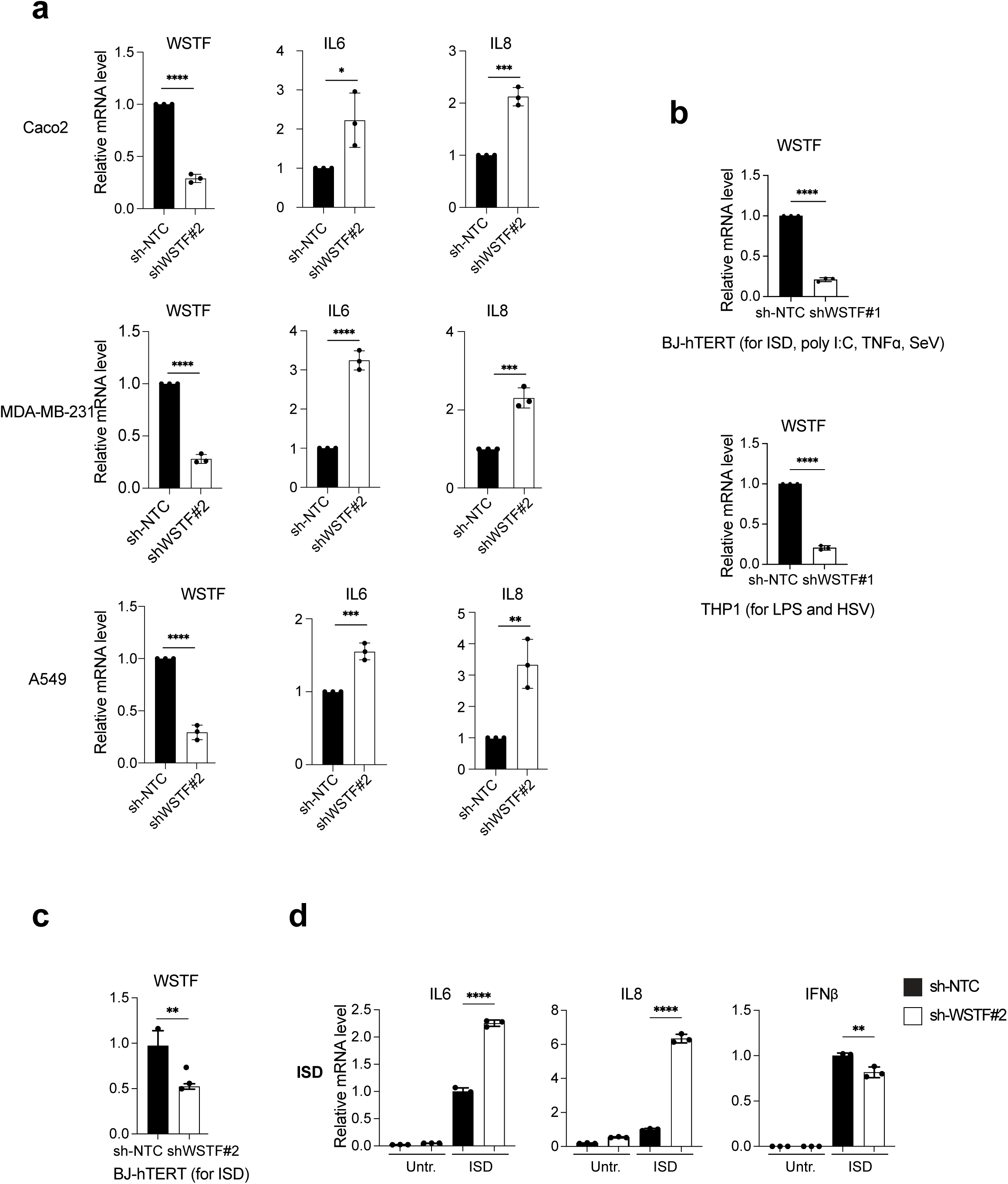
WSTF restrains the transcription of pro-inflammatory genes in cancer and other conditions. **a**, Related to Fig. 7a, RT-qPCR analyses of IL6, IL8 and WSTF in sh-NTC and sh-WSTF #2 cancer cells, including CaO.2, MDA-MB-231, and A549. Results were normalized to Lamin A/C and presented as mean values with s.d.; n=3; n.s. non-significant; * P < 0.05; ** P < 0.01; *** P < 0.001; **** P < 0.0001; unpaired two-tailed Student’s t-test. **b**, Related to Fig. 7c and d, RT-qPCR analyses of WSTF in sh-NTC and sh-WSTF#1 BJ-hTERT cells for ISD, poly I:C, TNFɑ and SeV, and THP1 cells for LPS and HSV-1. Results were normalized to Lamin A/C and presented as mean values with s.d.; n=3; n.s. non-significant; **** P < 0.0001; unpaired two-tailed Student’s t-test. **c,** RT-qPCR analyses of WSTF in sh-NTC and sh-WSTF #2 BJ-hTERT cells for ISD stimulation. Results were normalized to Lamin A/C and presented as mean values with s.d.; n=3; ** P < 0.01; unpaired two-tailed Student’s t-test. **d,** RT-qPCR analyses of IL6, IL8 and IFNβ in sh-NTC and sh-WSTF #2 BJ-hTERT cells transfected with ISD. Results were normalized to Lamin A/C and presented as mean values with s.d.; n=3; ** P < 0.01; **** P < 0.0001; unpaired two-tailed Student’s t-test.

**Extended data figure 10.**
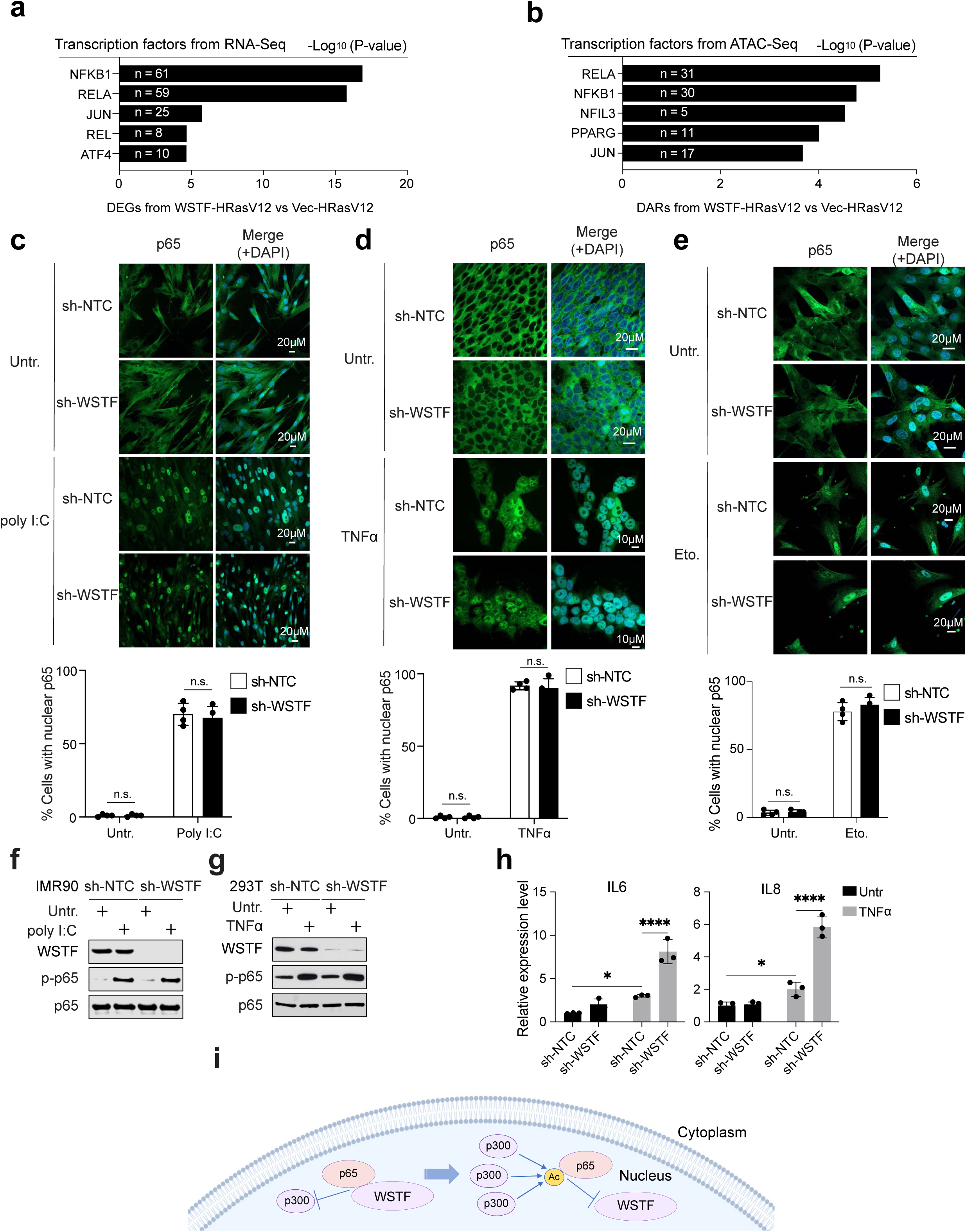
WSTF inhibits p65-mediated pro-inflammatory gene expression. **a**, **b**, Transcription factor prediction from RNA-seq using DEGs and ATAC-seq from DARs that are downregulated by WSTF in HRasV12-induced senescence. **c**-**e**, sh-NTC and sh-WSTF IMR90 (**c** and **e**) or HEK293T (**d**) were left untreated or treated with poly I:C 30 ug/ml for 1.5h, TNFα 20 ng/ml for 30min, or etoposide-induced senescence, then stained with a p65 antibody, followed by imaging under a confocal microscopy. Representative images are shown. Bar graphs showing the quantification of the percentage of cells with nuclear p65. Results are from four randomly selected fields with over 200 cells, presented as mean with s.d.; n.s.: non-significant. **f**, IMR90 cells were stably expressed with sh-NTC or sh-WSTF, and were treated with or without poly I:C, then analyzed by western blotting using indicated antibodies. **g**, 293T cells were stably expressed with sh-NTC or sh-WSTF, and treated with or without TNFα, then analyzed by western blotting using indicated antibodies. **h,** RT-qPCR analyses of IL6 and IL8 in sh-NTC and sh-WSTF 293T cells upon TNFα treatment. Results were normalized to Lamin A/C and presented as mean values with s.d.; n=3; n.s. non-significant; * P < 0.05; **** P < 0.0001; one-way ANOVA coupled with Tukey’s post hoc test. **i**, Schematic illustration of WSTF interaction with non-acetylated p65. Upon p65 translocation to the nucleus, WSTF binds to non-acetylated p65 and inhibits p300 binding to p65. Upon further stimulation, p300 acetylates p65, leading to dissociation between WSTF and p65, promoting p65 transcriptional activation of pro-inflammatory genes.

**Extended data figure 11.**
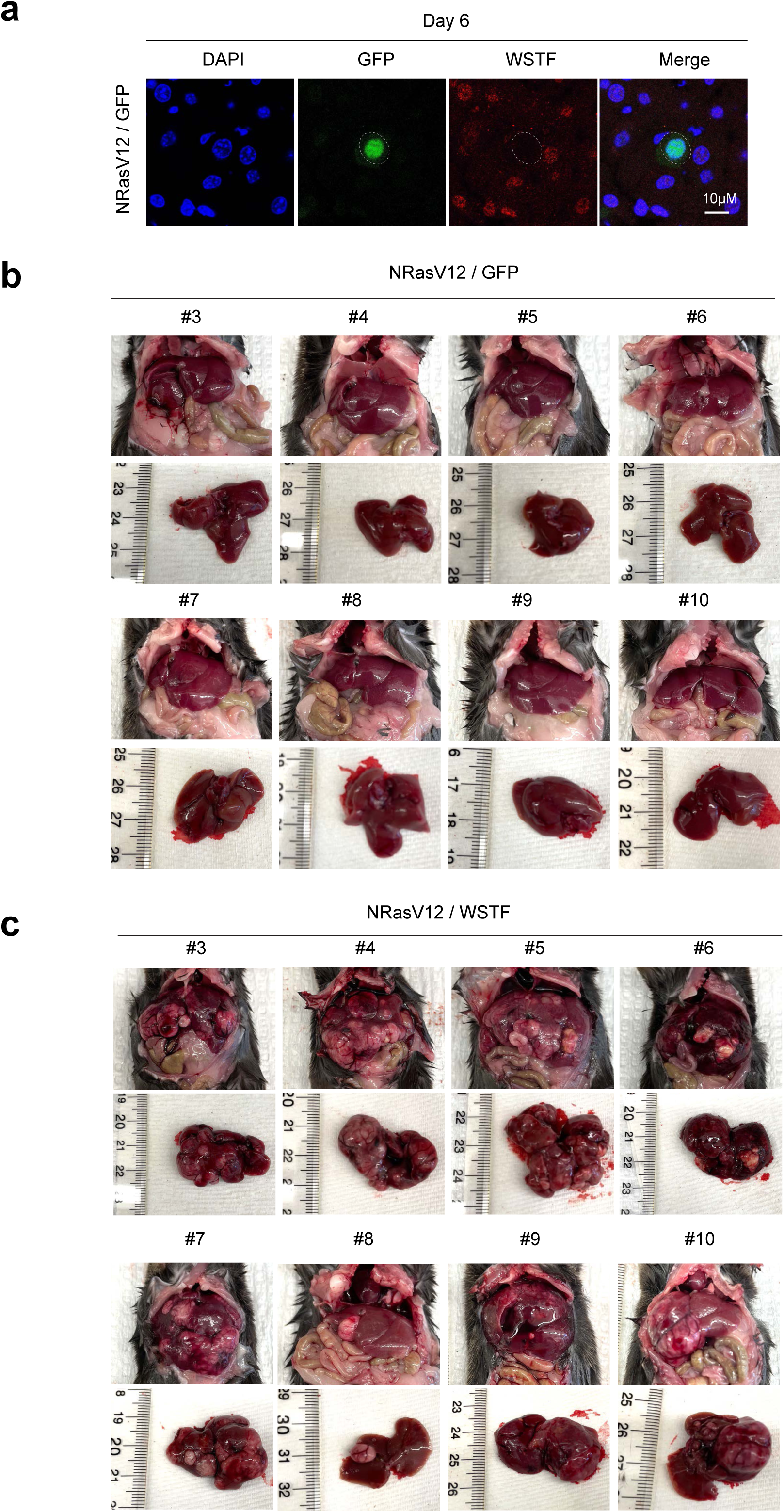
WSTF inhibits immuno-surveillance of NRasV12 in mouse liver. **a**, Liver sections 6 days post injection were stained with GFP and WSTF antibodies, and then analyzed by a confocal microscopy. Scale bars are shown. Note the GFP-positive cell showed reduced WSTF. **b** and **c**, Related to Fig. 8l and m, Images showing liver from NRasV12/GFP and NRasV12/WSTF groups 6 months post injection. Note while the GFP group did not develop liver tumors, all mice in the WSTF group developed liver tumors.

